# Unifying spatial and episodic representations in the hippocampus through flexible memory use

**DOI:** 10.1101/2025.09.21.677534

**Authors:** Xiangshuai Zeng, Jon Recalde, Laurenz Wiskott, Sen Cheng

## Abstract

A key question in neuroscience is why the hippocampus is essential for episodic memory in humans, while dominantly exhibiting spatial representations in a number of other species. Some accounts suggest that spatial representation is the primary hippocampal function. Here, we propose that the primary function is storing and retrieving episodic memories, and spatial representations emerge due to this memory function. To demonstrate this, we adopt a computational model that autonomously learns to store information in memory and retrieve it to solve a variety of tasks. In memory tasks, the model develops associations and categorical representations akin to concept cells. In navigation tasks, the model forms representations of the spatial structure, performs geometric computations, and even learns representations of unique events similar to recently discovered barcodes. Our model predicts that the hippocampus represents any task-relevant variable, if the animal learns the task, suggesting that space is not special for the hippocampus.

## 1 Introduction

Two landmark discoveries about the mammalian hippocampus led to the development of distinct research fields. Firstly, the discovery of memory deficits in patient H.M. (Scoville and Milner, 1957) led to theories emphasizing the hippocampus’ role in long-term memory. However, debates persist on whether the hippocampus should be viewed as an auto-associative (Cheu et al., 2012) or hetero-associative (de Camargo et al., 2018; Cheng, 2013) memory system. Secondly, the characterization of place cells in rodents (O’Keefe, 1976) led to theories emphasizing the hippocampus’ role in spatial navigation and investigations of spatial neural representations. One dominant theory posits that the hippocampal formation encodes a cognitive map of the environment (O’Keefe and Nadel, 1978; Jacobs, 2003; Jeffery and Burgess, 2006; McNaughton et al., 2006), which facilitates goal-directed navigation. This view received further support by later discoveries of other types of spatially-tuned cells in nearby brain regions connected, e.g., head direction (HD) cells in the subiculum (Taube et al., 1990) and grid cells in the entorhinal cortex (Hafting et al., 2005).

Despite significant progress in both fields and vigorous debates, few attempts have been made to reconcile these two views of hippocampal function in a computational model. Navigation models often neglect the presence of non-spatial cells in the hippocampus, and memory models typically do not incorporate the computational roles of spatially tuned cells. Attempts at reconciliation have taken two views. The first view is that spatial representations are key for memory (O’Keefe and Nadel, 1978; Chandra et al., 2025). Chandra et al. (2025) propose that a spatial scaffold, including grid and place cells, facilitates associative memory via hetero-associations between sensory inputs and the place cell module, and episodic memory via sequences generated by the recurrent grid cell module. The second view is that there is a common element of spatial navigation and episodic memory (Eichenbaum et al., 1999; Eichenbaum, 2017; Lisman et al., 2017), e.g., the “Tolman-Eichenbaum Machine” predicts the next sensory input by combining capabilities of memory storage and path integration (Whittington et al., 2020).

Here, we propose a third possibility: the primary purpose of the hippocampus is to store episodic memories, and spatial representations emerge due to this memory function. In this study, we use the term “episodic memory” in an operational sense, referring to the ability to encode information from a single experience and retrieve it later to guide behavior. This definition emphasizes one-shot encoding and flexible retrieval, rather than an explicit representation of temporal context (Howard and Kahana, 2002) or phenomenological aspects of recollection. To test this hypothesis we developed a computational model based on the fast weight memory (FWM) model (Schlag et al., 2021; Irie et al., 2021), a memory-augmented neural network (MANN), which allows flexible usage of a long-term memory (LTM) store. This model overcomes two major limitations of current hippocampal memory models: Firstly, they restrict representations to predefined formats and secondly, prescribe when and what information to store in memory. We study this model in a variety of pure memory and spatial learning tasks. We find that, in some tasks, the model learns to store information in LTM that we intuitively deem useful for task performance, e.g., an unmarked goal location for spatial navigation or the next element for sequence prediction. However, in other cases, unexpected representations and computations emerge, e.g., the memory retrieval process performs complex geometric computations in a navigation task. The model learns to exploit the memory and computational capacity of the LTM in unexpected ways that improve overall task performance and can account for a range of empirical findings related to both navigation and memory in mammals.

### 2 Results

### 2.1 Model and Task Overview

To model flexible encoding and retrieval of one-time experiences in LTM (Fig. 1), we adapted the FWM model (Schlag et al., 2021; Irie et al., 2021) to solve two pure memory tasks (Fig. 1c, d) with supervised learning (Fig. 1e) and two spatial tasks (Fig. 1f, g) that require memory with reinforcement learning (Fig. 1h). We refer to the model as an “agent” in the spatial tasks as is common in RL studies.

**Figure 1.**
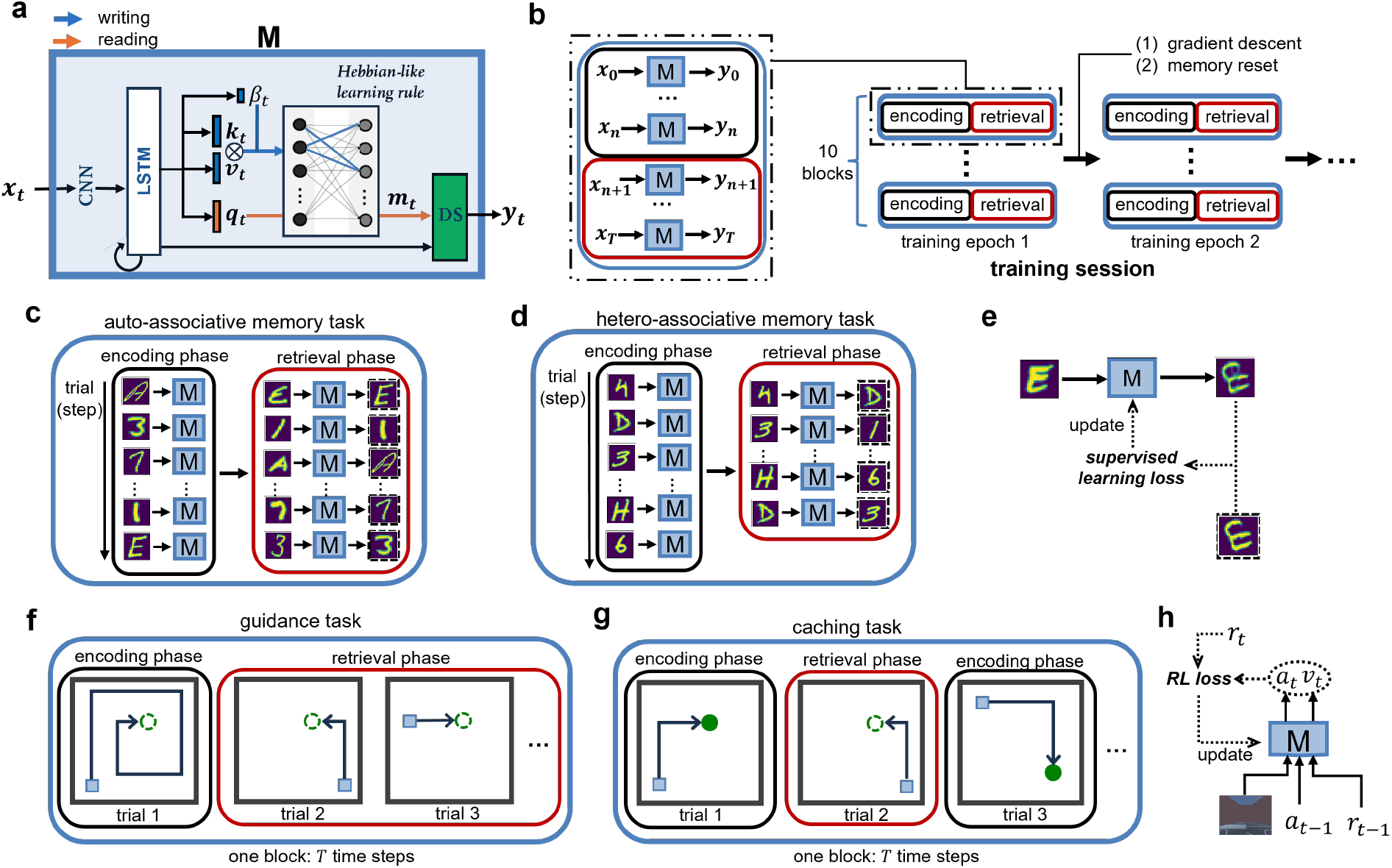
Illustration of the model and tasks. **a.** Model architecture. The entire model is referred to as “M” in other subplots. **x**_*t*_: input; **k**_*t*_: writing key; **v**_*t*_: writing value; *β*_*t*_: writing strength. **q**_*t*_: reading key; **m**_*t*_: reading value; **y**_*t*_: model output. The association between the writing key and value, multiplied with the writing strength, is written into the FWM matrix by a Hebbian-like learning rule (see Methods). The three blue and one orange rectangles represent trainable nonlinear layers. The network between the writing key and value layers represents the FWM matrix. The green rectangle represents the downstream network (DS) that is either a de-convolutional neural network in the memory tasks or a two-layer dense network in the spatial tasks. **b**. Task structure in the training and the test session. The encoding phase contains multiple trials in the memory tasks (c and d) and one trial in the spatial tasks (f and g). The retrieval phase contains multiple trials in both the memory tasks and the guidance task (f), but one trial in the caching task (g). In the training session, 10 parallel task blocks are deployed for each gradient update to increase the efficiency of data collection and training stability. In the test session, only one task block was used. Finally, between two training epochs: (1) gradients are computed and the model is updated; and (2) the FWM matrix and the LSTM hidden states are reset to zero. **c**. Auto-associative memory task. In the encoding phase, the model is presented a sequence of images of digits or uppercase letters. In the retrieval phase, the model has to output the exact encoded image based on a cue of the same class. **d**. Hetero-associative memory task. The encoding phase is the same as in the auto-associative task. During retrieval, the previous images are presented in random order and the model has to output the image that followed the current one in the encoded sequence. In both memory tasks, a “trial” refers to the presentation of one image to the model. Hence, one trial contains only one timestep. Between the encoding and retrieval phases, a blank image is presented. **e**. Supervised learning in memory tasks. The output image from the model and the ground-truth image are used to compute a supervised learning loss (see Methods). In the training session, the average of the loss over all 10 task blocks is computed to update the model via gradient descent. **f**. Spatial task termed “guidance”: The light blue square represents the moving agent, and the hollow, green circle an invisible goal. During encoding, the agent explores the environment to locate the invisible goal, triggering a strong, positive reward. The agent is then teleported to a new start location and can receive the reward at the same location repeatedly by returning to the goal location. The goal location changes after the block ends, which occurs after a fixed number of timesteps. **g**. Caching task. It is similar to the guidance task with two major differences: The goal is visible during encoding (solid, green circle) and the retrieval phase ends immediately after the agent reaches the goal location, after which a new encoding phase starts with a different goal location. For both spatial tasks, a “trial” refers to the time period between teleportation to a new location and the agent reaching the goal. Hence, one trial usually consists of multiple timesteps. **h**. Reinforcement learning in the spatial tasks. Based on visual input, previous action *a*_*t*−1_ and previous reward *r*_*t*−1_, the model generates the next action *a*_*t*_ and the state value *v*_*t*_ as outputs. An RL loss can be computed by using the output action and value together with the external reward signal. In actual training, an average of the RL loss is computed by using the data over all timesteps and task blocks.

At each timestep, external input **x**_*t*_ (gray or RGB images) to the model is pre-processed by a convolutional neural network (CNN) and passed to a long short-term memory (LSTM) model whose output is projected to different memory heads to generate four key components (Fig. 1a): the writing strength (*β*_*t*_), the writing key (**k**_*t*_), the writing value (**v**_*t*_), and the reading key (**q**_*t*_). Information is written (blue arrow) into the FWM matrix through a Hebbian-like learning rule (see Methods). The reading key is multiplied with the memory matrix (orange arrows) to yield the reading value (**m**_*t*_), which is further combined with the LSTM output and processed by the downstream (DS) network (green rectangle) to obtain the final output (**y**_*t*_). The CNN, the LSTM, the memory heads (nonlinear layers) and the downstream network are all trained “slowly” using gradient descent, while the FWM matrix is updated “fast” in every timestep using matrix operations. This way, the model learns to flexibly generate memory components that are useful for solving the task.

There is no pre-defined strategy for using the LTM, and the model has to learn autonomously what and when to perform memory writing and reading to optimize its task performance. Throughout the paper, we extensively analyze the writing and reading processes to understand how the model mechanistically solves a task.

All tasks are organized hierarchically (Fig. 1b): A task block includes an encoding phase, in which information is available once, and a retrieval phase, in which the previous information is needed to solve the task. An exception to this structure is the caching task (Fig. 1g) in which encoding and retrieval phases are repeated. Depending on the task, each phase consists of one or multiple trials that include one or multiple timesteps.

### 2.2 Auto- and Hetero-Associative Memory

Memory models are classified as auto-associative or hetero-associative, and it is debated what type of memory the hippocampus is (Rolls, 2010, 2013; Fortin et al., 2002; Cheng, 2013). We demonstrate that the same network can solve both types of memory tasks. In an auto-associative memory task, the model has to reconstruct the exact encoded image based on a similar cue (Fig. 1c). In a hetero-associative memory task, the model has to retrieve the next item in a memorized sequence when presented with a list item (Fig. 1d). As stimuli, we chose the extended MNIST dataset containing handwritten digits and Roman letters (Cohen et al., 2017). To reduce visual ambiguity (e.g., between “1” and “I”), we restricted our dataset to the first 18 classes (digits 0–9 and letters A–H).

The model successfully learned the auto-associative memory task (Fig. 2a). Writing to LTM occurred only during encoding, indicated by the vanished writing strength during retrieval. To examine what was written to and read from the memory matrix, we fed the downstream network with the writing and reading value vectors to reconstruct images (see Method 4.3). The input images in the encoding phase were reconstructed from both the writing values during encoding and the reading values during retrieval (Fig. 2a, second row in both upper and lower parts), indicating memory storage during encoding and correct associative recall during retrieval.

**Figure 2.**
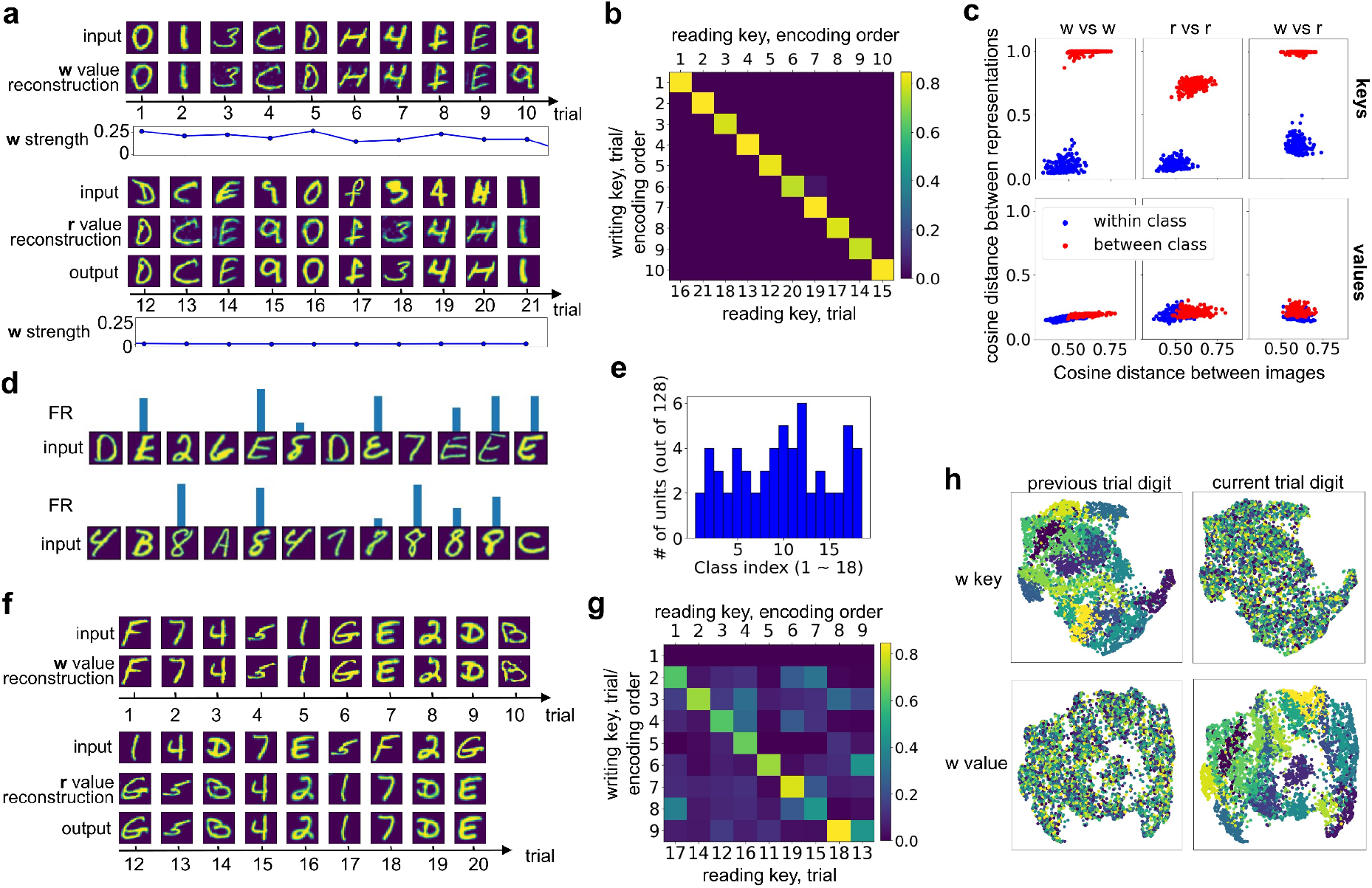
Task-specific representations in auto-associative and hetero-associative tasks. The letters”w” and “r” indicate writing and reading, respectively. **a**. Sample test block in the auto-associative task. The upper and bottom half corresponds to the encoding and retrieval phase, respectively. The buffering trial (no. 11) and the output in the encoding phase are not shown because they are not relevant to the analysis. The reconstructed images from the writing and reading values (without concatenating with LSTM output) show that the model wrote the complete sensory details of the input images to memory and later retrieved them. Note that the output images are generated by the concatenation of the LSTM output and the reading value. **b**. Cosine similarity between the writing keys during encoding (rows) and the reading keys during retrieval (columns). Columns are re-arranged based on the encoding order, shown at the top. **c**. Distances (1-cosine similarity) between memory items reveal strong categorical representations of keys, but not of values. For each point, 20 samples of one class randomly drawn from the testing data were compared to 20 samples from the same class (blue points) or from the another class (red points). The pair-wise cosine distances between the corresponding input images and between the memory items were computed and averaged to form one point. The headers and row labels indicate which memory items were compared. The different ranges of distances between images in the case of “w vs r” stem from the fact that the cue images used in the encoding and retrieval phases are similar within a phase, but different across the phases. **d**. Two examples of concept units in the writing keys that are selective for one class. FR: firing rate, the activity level of an artificial neuron. **e**. Summary of selectivity. On average, 60 out of 128 units in the writing key during encoding exhibited class-selective responses, while the rest remained silent throughout the test session. Each class activated at least two units. **f**. Sample test block in the hetero-associative task. Upper and bottom half corresponds to the encoding and retrieval phase, respectively. **g**. Cosine distance between writing keys during encoding (rows) and reading keys during retrieval (columns). Columns are re-arranged based on the encoding order, shown at the top. A writing key in trial *t* has high similarity with a reading key in trial *t* − 1. **h**. 2D U-map projections reveal that during encoding, writing keys represent the image class from the previous trial, while the writing values represent the image class from the current trial.

As required for task performance, writing keys during encoding were aligned with reading keys during retrieval (Fig. 2b). Writing and reading keys were very similar for images of the same class and diverged for different classes, revealing a categorical representation of the images (Fig. 2c, top), which was not present at the image level. By contrast, value vectors did not exhibit categorical representations (Fig. 2c, bottom) and their variance was even smaller than that of the input images, because of the smaller dimensionality (128 vs 784). Principal component analysis (PCA) confirmed that key vectors cluster by class, whereas value vectors do not (Supplementary Fig. 1), In other words, the model discovered that solving the auto-associative task requires the separation of categorical representations/ semantics (key vectors) and overlapping sensory content (value vectors).

At the single-unit level, many units in the writing key were selective to individual image classes (Fig. 2d, e), resembling concept cells found in the human hippocampus (Quiroga et al., 2005). A similar pattern was found in reading key units, some of which preserved class identity, but showed less sparsity (Supplementary Fig. 1a, b, unit 0). In contrast, units in the value vectors either responded non-selectively or remained silent (Supplementary Fig. 1 c, d), matching the distributed pattern in the population-level representations.

The same model also learned the hetero-associative memory task, i.e., when cue with an image during retrieval, the model generated the next image in the encoding sequence (Fig. 2f). The model learned to store the hetero-association between successive images by representing the *previous* item as the writing key and the *current* item as the writing value during encoding. This is suggested by two pieces of evidence. Firstly, similarity was highest between reading keys at encoding order *i* and writing keys at encoding order *i* + 1 (Fig. 2g). Secondly, in a 2D UMAP projection (McInnes et al., 2020), writing keys clustered based on the previous image, and writing values clustered based on the current image (Fig. 2h).

The model could retrieve the entire sequence, when cued with the first image and iteratively using the output as the next retrieval cue, although the results were not always perfect (Supplementary Fig. 2a). When comparing the self-generated sequences with those, where external cues were provided for each retrieval, two notable patterns emerged (Supplementary Fig. 2b). Firstly, construction errors accumulated with the position in the sequence in the self-generated sequence, but not in the cue-driven sequence. Secondly, for both retrieval modes, reconstruction errors increased with the length of the stored sequence due to increased memory interference.

Finally, the model generalized to the unseen 18 image classes in both tasks (Supplementary Fig. 2c-f). Performance was better in the hetero-associative task, probably due to the underlying geometry of the learned representations. The hetero-associative task leads to a more continuous, less separated key space (Fig. 2h), supporting better generalization. In contrast, in the auto-associative task, writing keys form orthogonal, categorical clusters (Supplementary Fig. 1a), limiting generalization to unseen classes.

In summary, a model with the same network architecture solved both auto- and hetero-associative memory tasks by adapting what and when to encode and retrieve, highlighting the model’s computational versatility.

### 2.3 Attractor Dynamics in Emergent Spatial Representations

CA3 has been hypothesized to be an auto-associative memory network, with attractor dynamics proposed as one possible mechanism. This hypothesis was tested in spatial settings by morphing the environment between a square and a circle (Supplementary Fig. 3a-f) (Leutgeb et al., 2005; Wills et al., 2005; Colgin et al., 2010). The population vectors (PVs) of CA3 activity in the morph environment were compared to the endpoints of the sequence, i.e., the square and the circle maze. While Wills et al. (2005) observed an abrupt shift of similarity, consistent with a sharp transition between two attractors, Leutgeb et al. (2005) reported a gradual change. Colgin et al. (2010) suggested that these divergent findings were caused by different training conditions: a “single-location” condition, where rats alternated between the square and the circle maze, presented at the same location, and a “double-location” condition, where the square and circle were located at different positions and connected by a corridor (Supplementary Fig. 3g). When tested in the intermediate morph environments later, the single-location condition indeed induced gradual changes between representations (Fig. 3a, top-right), whereas the double-location condition produced abrupt shifts (Fig. 3a, bottom-right) (Colgin et al., 2010).

**Figure 3.**
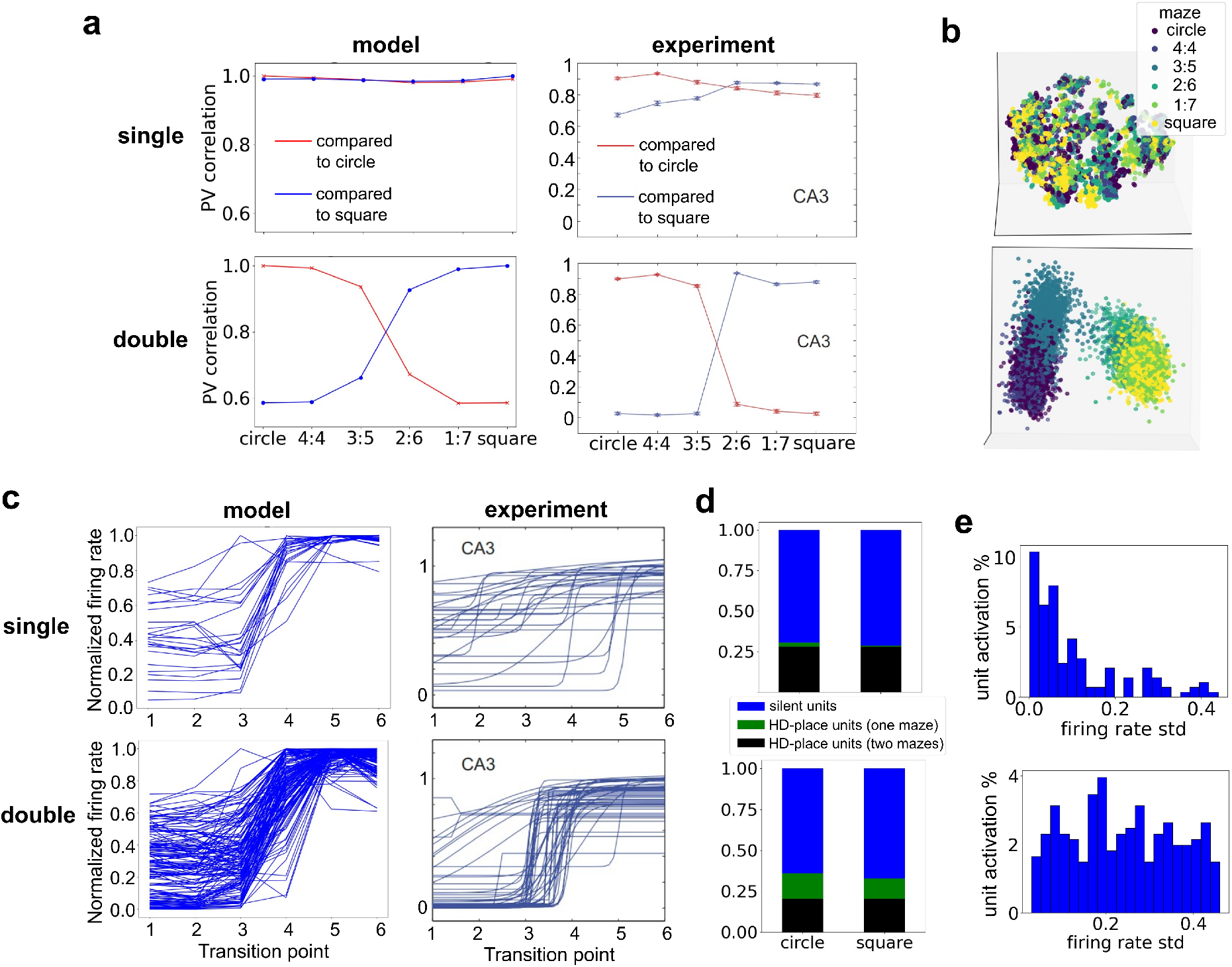
Accounting for attractor dynamics in the hippocampus revealed by morph experiments. The labels “single” and “double” refer to the single-location and double-location condition, respectively, “experiment” to the data reproduced from Colgin et al. (2010) with permission. **a**. Population vector correlations across the environment morph sequence, computed from the head direction (HD)-place units in the reading keys, averaged over 10 simulations (left) and place cells in the CA3 region of the hippocampus (right). **b**. 3D PCA projections of the reading keys during retrieval from a typical simulation. The color of the points indicates the maze shape. **c**. Remapping for individual units. The order of shapes of the mazes was reversed for rates that decreased across the sequence so that all rate changes could be plotted in increasing order. Note that remapping occurs only in a few units in the single-location condition. Left: Average firing rates of those units in the reading key that passed a sigmoid fit function (see Methods), selected from all 10 simulations. Right: Sigmoid fits for in-field firing rates for all place fields in CA3 that surpassed the 2-Hz threshold, selected from all animals of the corresponding group in the experiment. **d**. Proportion of silent units (blue) and HD-place units in the reading keys in the circle and square mazes under the two conditions, averaged over 10 simulations. HD-place units are further subdivided into those that are active in only one maze (green) and that are active in both mazes (black). **e**. The distribution of standard deviations (std) of the firing rates of units in the reading keys that are active in at least one maze across the six test mazes, pooled over 10 simulations. A condensed density around 0 std means that the firing rates are stable across the mazes.

We replicated these experiments in our model using a navigation task (called “guidance”), in which the agent/model has to search for a reward during encoding and return to this location repeatedly during retrieval to receive rewards (Fig. 1f). Since the reward location is unmarked, the agent is solely guided to the goal location by distal cues (Parra-Barrero et al., 2023)). To maximize the total reward, the agent has to remember the goal location and reach it efficiently from anywhere in the maze. To make the shape of the environment behaviorally significant for the agent, there were no step penalties in the corners of the square environment, reflecting rodents’ preference for corners (see Methods). The agent learned the task in both training conditions (Supplementary Fig. 4c). In the test session, the number of steps to reach the goal decreased rapidly after the first encounter (Supplementary Fig. 4a), confirming one-shot learning of the goal location. This behavior generalized to other environments in the morph sequence without additional training (Supplementary Fig. 4b).

The model learned spatial representations similar to experimentally recorded ones (Supplementary Fig. 5b, d). Focusing on the reading key in the retrieval phase when memory is most actively queried, single units are selective to a conjunction of the agent’s HD and location (Supplementary Fig. 5a, c). We return later to this issue of mixed representations, but note here that individual units are modulated by the agent’s position, similarly to place cells. The PV correlations of the reading key (see Methods) reproduced the experimentally observed differences between the two conditions (Fig. 3a). PCA projections revealed well separated clusters for circle-like and square-like mazes in the double-location condition (Fig. 3b, bottom), demonstrating two separated representations. By contrast, in the single-location condition (top), representations for the differently shaped environments were more continuous. This difference in representations aligns with the agent’s behavior: in the single-location condition, behavior was nearly homogeneous, despite consistently receiving less punishment at the corners of the square maze across training conditions, whereas in the double-location condition the agent spent significantly more time at the four corners of the three square-like mazes (Supplementary Fig. 4d).

Many of the units in the model exhibited global remapping, an abrupt change in activity patterns, near the midpoint of the morph sequence in the double-location condition (Supplementary Fig. 5c), but less so in the single-location condition (Supplementary Fig. 5a). Fitting sigmoid functions to in-field firing rates (Colgin et al., 2010), revealed that more HD-place units showed abrupt transitions in the double-location condition (39 of 71, 55%, Fig. 3c, bottom-left) than in the single-location condition (6 of 33, 18%, Fig. 3c, top-left), consistent with experimental results (Fig. 3c, right). In contrast to the experiments, where the transitions occurred at multiple points along the shape sequence (Fig. 3c, top-right), the transition of units in the model occurred mostly at the mid-point (Fig. 3c, top-left). Most HD-place units in the single-location condition are active in both square and circle maze (Fig. 3d, top), and had stable firing rates, since most of the standard deviations of the in-field firing rates were near zero (Fig. 3e, top).

In summary, our model exhibits representations that were previously interpreted as evidence for attractor dynamics in the hippocampus at both the single-unit and population level, without being an attractor network. Instead, the effects arose through learned adaptation to task demands, particularly the need to distinguish different environments, and act accordingly.

### 2.4 Barcode and Spatial Code: Linking Episodic Memory and Spatial Representations

Recently, Chettih et al. (2024) reported that CA1 neurons simultaneously encode information about space and episodic-like events, when chickadees moved around in a large arena while caching seeds in covered sites and later retrieving them. CA1 neurons exhibited two distinct neural signals: a smooth place code that reflected the bird’s location (Fig. 4a, bottom) and discrete, event-specific “barcodes” when food was cached (Fig. 4a, top) or retrieved (Fig. 4a, middle) in the same location. Importantly, both place cells and non-place cells participated in the barcodes, and the preferred location of a place cell could differ between the spatial code and a barcode.

**Figure 4.**
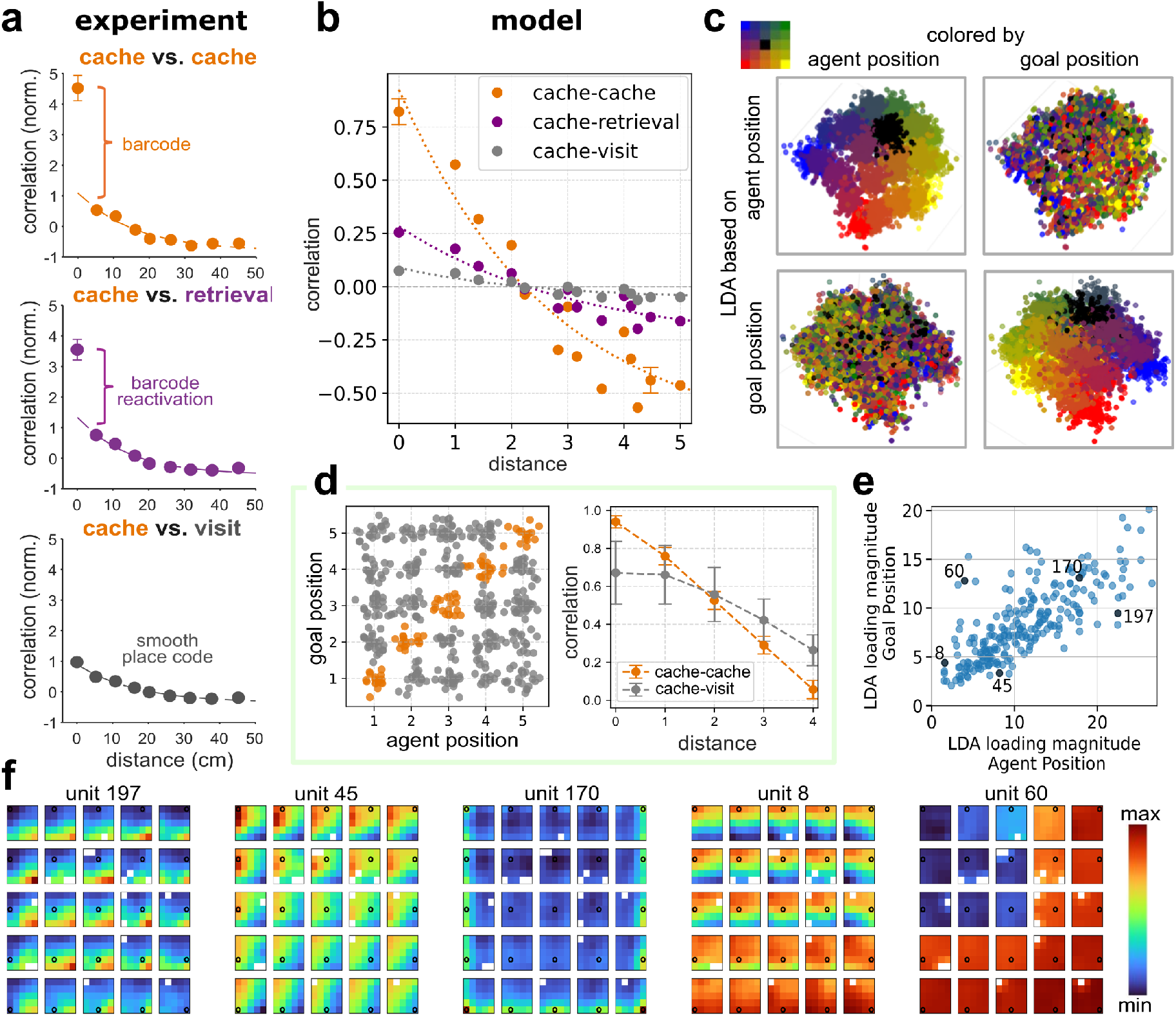
Linking spatial code with barcode representation of episodic memory. **a.** Population vector (PV) correlations for hippocampal CA1 as a function of spatial distance for three different behavioral comparisons (reproduced from Chettih et al., 2024, with permission). Strong correlations at 0 cm for cache–cache and cache–retrieval and steep drop for larger distances indicate an episode-specific barcode that reactivates when the bird returns to the same site. Cache–visit events, where visits lack retrieval intent, show a smooth decay with distance, consistent with place coding. **b**. PV correlations in our model for LSTM activity in the trained model. Cache–cache (orange) and cache–retrieval (purple) pairs show stronger similarity at shorter distances compared to cache–visit pairs (gray), replicating the structure observed in the experimental data. **c**. Linear discriminant analysis (LDA) of LSTM activity. PVs are projected onto the first two discriminant axes obtained from training on either agent position (top row) or goal position (bottom row) labels. Each plot is colored by agent position (left) or goal position (right). The clustering and topographic organization indicate that both variables are distinctly and independently encoded by the model. **d**. Simplified model used to isolate the role of spatial features. Left: PV activity for individual events: cache events (orange) tightly cluster along the diagonal, where agent and goal location coincide, while visit events (gray) are more distributed across the columns. Right: correlations between event pairs: cache–cache pairs show higher similarity at distance 0 and a sharper decrease with distance than cache–visit pairs. **e**. Goal and agent location coding in single LSTM units. Axes show the loading magnitude (norm of each unit’s loading vector) from two-component LDA trained with agent-position labels (x) or goal-position labels (y). Pearson correlation r=0.84, indicating that units coding for agent position also tend to code for goal. Black dots indicate the example units in panel f. **f**. Spatial activity maps for five representative LSTM units during retrieval. For each unit, the 25 maps represent the 25 possible goal locations (black circle) and show mean activity at different agent locations, scaled to the unit’s own activity range. Empty nodes are those that the agent didn’t pass through under the conditions.

To examine how barcoding could arise and exist in parallel with the spatial code, we trained our model on a simplified version of Chettih et al.’s task using reinforcement learning (Fig. 1g, Supplementary Fig. 6a,b). In the encoding phase, the agent has to navigate to a goal location marked by a white sphere (called “aiming”, Parra-Barrero et al., 2023), while in the retrieval phase it has to navigate to the same, now unmarked, location (guidance). In addition to the navigation actions, the agent has an *open site* action that allows it to cache food during encoding and retrieve food during retrieval (see Methods). The trained agent consistently cached and retrieved at the correct location, indicating one-shot memory encoding and retrieval. We examined the correlations of LSTM outputs between different events, since the LSTM outputs project to both memory-writing and memory-reading pathways and provide a common representational space in which cache, retrieval, and visit events can be directly compared. We found that the correlations were much higher between cache events at the same location than at nearby ones (Fig. 4b). There is also a clear difference between cache–cache, cache–retrieval, and cache–visit correlations in the model.

**Figure 6.**
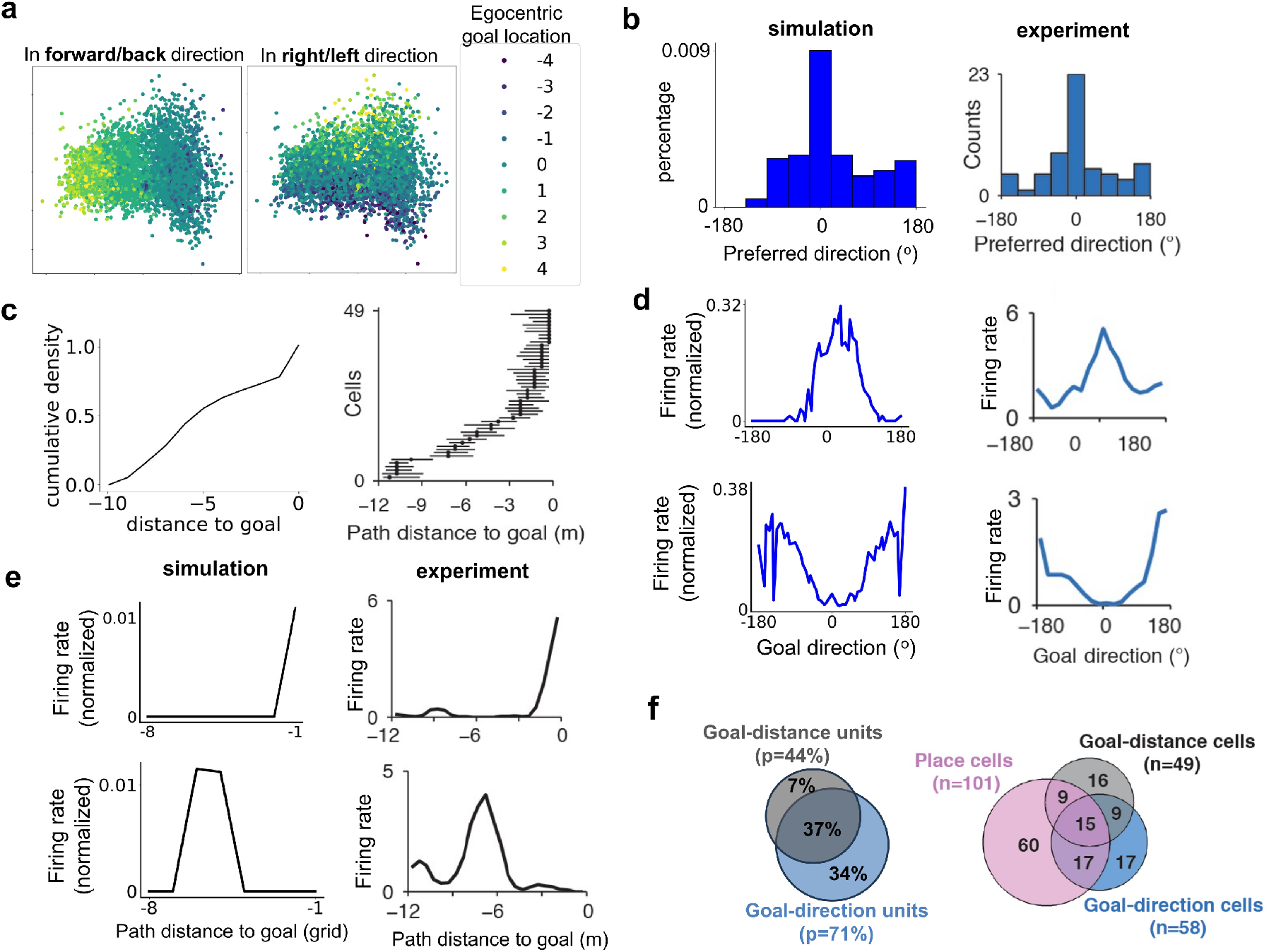
Egocentric goal representations in model account for data from bat hippocampus. In all panels, simulations are shown on the left and bat data (reproduced from Sarel et al., 2017, by permission) on the right. **a**. PCA projections of the reading values, colored by egocentric goal location in the forward/backward direction (left) and in the in the right/left direction (right). Positive values in the labeling refer to the goal being in the front/right of the agent. **b**. Histogram of preferred direction relative to goal. **c**. Cumulative distributions of preferred distance to goal. **d**. Examples of tuning curves of the goal-direction unit. **e**. Examples of tuning curves of the goal-distance unit. **f**. Fraction of goal-direction units, the goal-distance units and place cells.

To better understand the internal representations and what gives rise to the barcode, we performed linear discriminant analysis (LDA) on the LSTM population vectors based on either agent or goal position. Projections revealed that both variables are encoded along distinct axes in the latent space (Fig. 4c). This mixed representation suggests that barcodes may arise from unique combinations of several variables that coincide for individual events, but are not necessarily all identical when the agent merely passes the same location. To test this hypothesis, we built a simplified model using synthetic, joint representations of two 1D variables: agent and goal positions (Fig. 4d, left) (see Methods). Cache events (orange) cluster tightly along the diagonal where agent and goal positions coincide and therefore generate high correlations between population vectors. As the distance between cache sites increases, both dimensions change, leading to a steep decrease in correlation (Fig. 4d, right). By contrast, visit events, where only agent locations coincide and goal positions vary, are more distributed across a column (Fig. 4d, left, gray). This spread in representational space leads to greater variability across events and lower average correlations (Fig. 4d, right). Since the drop-off of the correlation with distance is smoother in the simplified 1D model than in our network model (Fig. 4b), we hypothesize that the even steeper drop-off in the experiment (Fig. 4a) stems from a higher dimensional representational space.

We next examined how individual units respond. We found a linear relation (Pearson *r* = 0.84, *p* < 10^−74^) between the Euclidean-norm of each neuron’s loading vector within the LDA subspace for the agent position and that for the goal position (Fig. 4e), suggesting that neurons contributed similarly to both dimensions, but to varying degrees. Decoding accuracy of agent and goal locations remained high until more than half the neurons were silenced (Supplementary Fig. 6d), suggesting that the barcode is distributed across the population rather than carried by individual units. This conclusion is further supported by heterogeneous response properties of individual units with varying levels of goal-modulation (Fig. 4f). Some units behave like a classical place cell (unit 197); modulate their place field activity depending on the goal location (unit 45); shift their field toward the current goal position (unit 170); indicate whether the agent is located above or below the goal (unit 8); or fire almost exclusively when the remembered goal occupies one specific region, independent of the agent position (unit 60).

In conclusion, we suggest that barcodes are a result of highly mixed selectivity in hippocampal neurons that simultaneously code for several latent variables. The more variables match between two time points, the higher the correlation between the corresponding hippocampal representations. When calculating spatial correlations we consider only two positional dimensions, ignoring other variables in the high-dimensional space, which lowers the correlations as compared to event-event correlations.

### 2.5 Emergence of Spatial Computations in Memory Network Explains Goal-Vector Cells in Bats

One of the keys to understanding spatial navigation behavior are the computations performed over the spatial representations (Vijayabaskaran et al., 2025). We therefore trained the agent on the guidance task (Fig. 1f) only in a single environment and analyzed the trained model. We found that the writing strength *β*_*t*_, which gates writing into the FWM matrix, exhibited a large peak every time the agent reached the goal (Fig. 5a). This peak was especially high the first time the agent reached the goal, denoted as *t* = *g*, suggesting that the model encodes information about the goal location. Low writing strength observed at other times helps avoid memory interference. PCA on the writing keys **k**_*t*_ at *t* = *g* from all trials in the test session reveal clear representations of the goal locations (Fig. 5c). This pattern does not emerge for writing values **v**_*g*_ (Fig. 5d), raising the question how easily location information can be extracted from LTM. In fact, PCA on the flattened FWM matrix, **F**_*t*_, recovers representations of goal locations (Fig. 5e). In the retrieval phase, we expected that the agent would retrieve the goal information from the FWM memory, so that the downstream network can compute the action based on the goal location and the current location supplied by the LSTM. To our surprise, we found that the reading keys **q**_*t*_ formed four banana-shaped PCA clusters, each associated with one HD (N, E, S, W) (Fig. 5f, g). In pairs of opposing directions (e.g., N/S or E/W), projections emphasized the agent’s coordinates along the axis aligned with the pair of corresponding HDs (Fig. 5h). Surprisingly, the reading value **m**_*t*_ reflected the action the agent will take (Fig. 5i), not the goal location.

**Figure 5.**
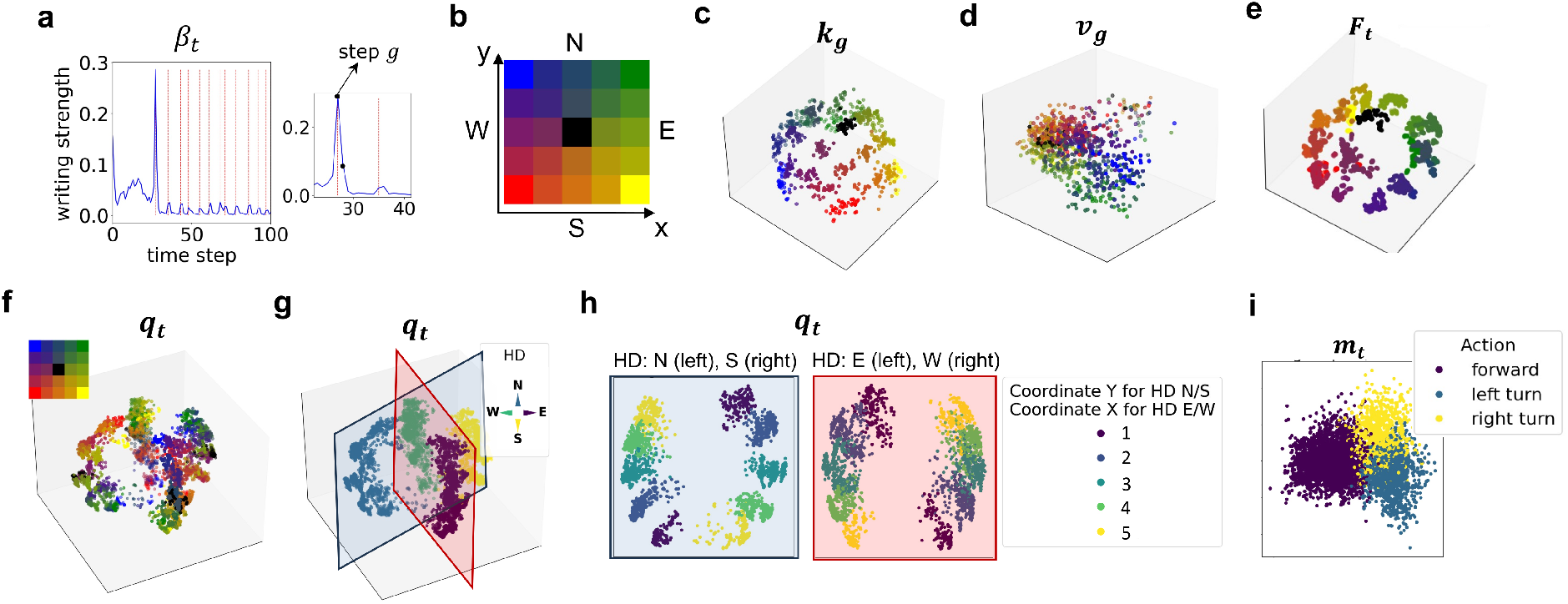
Spatial representations emerge in the PCA projections of the memory representations. All data are from the testing session only. **a**. Writing strength in an example trial. Red lines indicate the steps where the goal is reached by the agent. The inset on the right panel shows the first two times the agent reaches the goal. The time span between the first and second goal encounters is the first retrieval trial. The values at the peak (*t* = *g*) and right after the peak (*t* = *g* + 1) are marked by small, black points. **b**. The color scheme used for indicating the grid location in the environment, with the X, Y coordinate axis and the four head directions (North, South, East, West). **c**. 3D PCA projections of writing key **k**_*g*_ at step *t* = *g*. Points are colored according to goal location with the indicated color scheme in panel b. **d**. Same as in c, but for the writing value **v**_*g*_. **e**. PCA projections of the flattened fast weight memory matrix **F**_*t*_ during the entire retrieval phase (*t* > *g*), colored by the goal location. **f**. PCA projections of the reading keys **q**_*t*_ during retrieval (*t* > *g*), colored by the agent’s location. Color scheme as in panel b. **g**. Same projection as in panel f, colored by the head direction (HD) of the agent. **h**. Left: projections of the reading keys filtered for HD=north or HD=south, colored by the Y coordinate of the agent. Right: projections of the reading keys filtered for HD=east or HD=west, colored by the X coordinate of the agent. **i**. PCA projections of the reading values **m**_*t*_ during retrieval, colored by the action that the agent takes.

How does this representation of action arise? A detailed explanation is given in the Supplementary Text 6.1. Briefly, this unexpected representation stems from the geometric structures of the reading and writing keys in a common PCA space and the writing value at step *g* + 1. Specifically, **k**_*g*_ and **q**_*t*_ lie on the same spherical structure in 3D space, offset by 90° (Supplementary Fig. 7b), so that their dot product encodes the goal’s distance along the agent’s HD. Meanwhile, the writing value **v**_*g*+1_ encodes the goal’s relative position perpendicular to this direction. Combining all these, we found that the memory readout actually represents the ego-centric goal position (Fig. 6a), and the agent moves forward when the goal is ahead and turns left or right when it is behind, generating the observed action representations. Two lines of evidence indicate that the emergent representations and computations have a causal role in the goal-directed behavior. Firstly, their emergence during learning is correlated with improvements in behavior (Supplementary Fig. 8). Secondly, they generalize to nearby locations that were not explicitly trained as goal locations (Supplementary Fig. 10).

We next examined whether the geometric structure also manifests at the single-unit level. In the writing keys **k**_*g*_, some units selectively respond to the allocentric goal location, independent of the agent’s HD (Supplementary Fig. 9a). Units in reading keys **q**_*t*_ encode both HD and spatial location (Supplementary Fig. 9b). The reading values **m**_*t*_ contain units that encode the egocentric location of the goal (Supplementary Fig. 9c), matching the geometric analysis in the PCA space (Fig. 6a).

These findings parallel empirical observations in CA1 of freely flying bats during goal-directed navigation (Sarel et al., 2017). Some cells were tuned to the angle between the bat HD and the vector to the goal (Fig. 6d, right), or the Euclidean distance to the goal (Fig. 6e, right). Remarkably, our model reproduces both tuning types in reading value units (Fig. 6d, left; Fig. 6e, left). The distributions of the units’ tuning properties are consistent with biological data: Goal-direction tuning spans the full angular range from 180° to −180°, with a notable peak near 0° (Fig. 6b), i.e., when the goal lies directly ahead. Similarly, goal-distance tuning covers all distances, although short distances are not as over-represented as in the experiment (Fig. 6c). Furthermore, we found units exhibiting joint tuning to goal-direction and distance (Fig. 6f, left), consistent with compound “goal-vector” cells reported in experiments. Unlike the experiment, the model did not generate classic place cell-like activity in the reading values, suggesting that the experimental recordings were obtained from cells corresponding to both reading keys and values in our model.

In summary, a complex geometric computation arises from the operation of the FWM memory matrix and the geometric structure of representations that the model learns for the reading and writing keys and the writing value. The result is an egocentric goal representation in the reading value that appears at the population level and in the tuning properties of individual units, consistent with hippocampal recordings.

### 2.6 Unconstrained Spatial Movement Leads to Reduced Directionality in Place Fields

Hippocampal recordings in open fields show that, while firing rates are modulated by HD, place field locations remain relatively stable (Muller et al., 1994; Rubin et al., 2014; Jercog et al., 2019). However, the degree of directional modulation can depend on sensory and behavioral context, and place field selectivity has been shown to be shaped by the availability and integration of multi-sensory cues (Ravassard et al., 2013) In the simulations above, rate maps of reading key units rotate with the HD (Fig. 7a-e, Supplementary Fig. 5a and c). Notably, when rodent movement is restricted in experiments, e.g., on linear tracks (McNaughton et al., 1983; Battaglia et al., 2004) or by task demands (Markus et al., 1995), place field positions change with HD. So, the strong HD-modulation of place fields in our model may arise from a key restriction: the agent moves on a discretized grid with four HDs.

**Figure 7.**
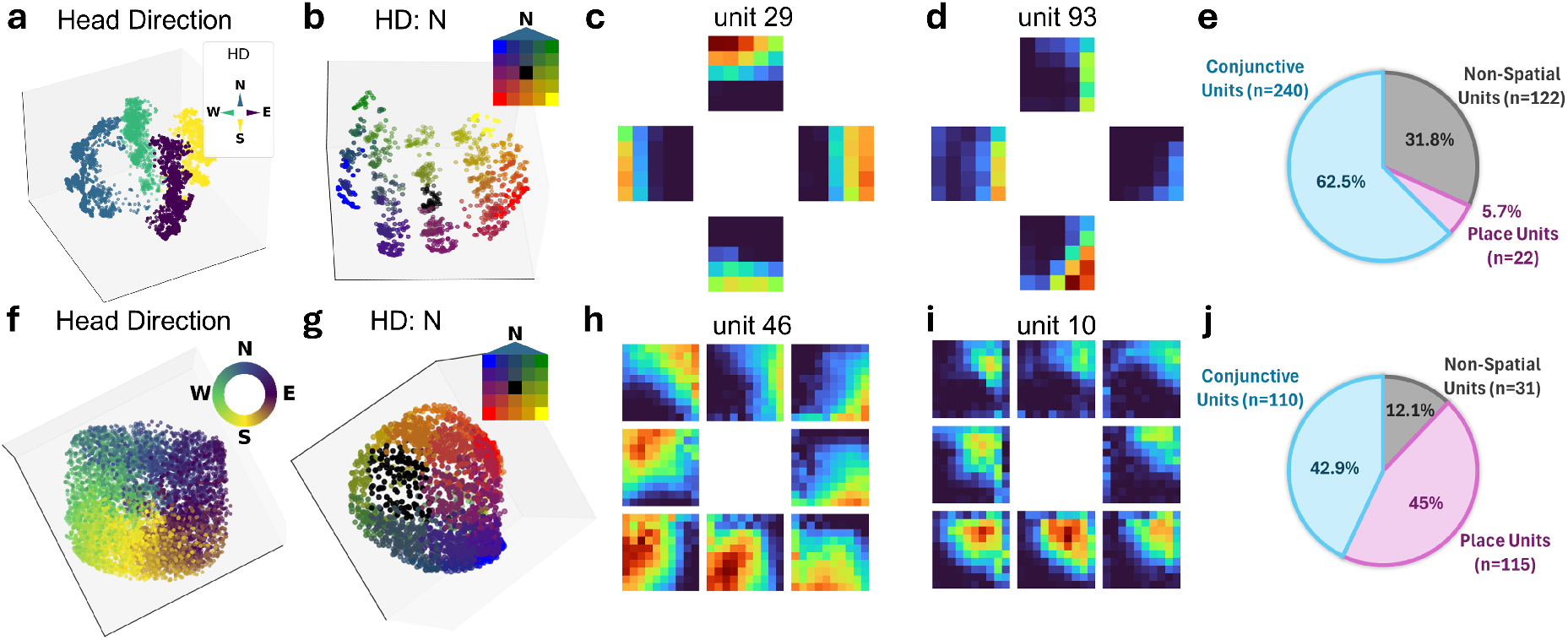
Simulations in continuous state space yields head-direction (HD) invariant spatial representations. a–e. In a discrete 5×5 grid environment, representations of agent’s location rotate with HD. **f–j**. In continuous state space, spatial representation of some units are HD-invariant. **a, f**. PCA representations of reading keys colored by HD. **b, g**. Same PCA representations filtered for when the agent is facing north (–22.5° to 22.5°), colored by the agent position. **c, h**. Example of a conjunctive (HD×place) unit: firing rate maps shown separately for different HD. For the state space, we binned HD into 45° sectors, allowing us to add the four inter-cardinal directions (NE, SE, SW, NW) in panels h-i. **d, i**. Example of a place unit: firing rate maps remain consistent across all HD. **e, j**. Fractions of conjunctive, place-only, and non-spatial units.

To test this hypothesis, we trained the agent on the guidance task in a continuous state space (see Methods). As before, reading keys encoded both HD and location (Fig. 7f, g) and the spherical structure in PCA space is consistent with the four banana-shaped clusters in the discrete case (Fig. 7a), suggesting that the computation of the egocentric goal location discussed above for the discrete case is preserved in the continuous case. Importantly, some single units now exhibit stable place fields across HDs (Fig. 7i, j). Nevertheless, other units were clearly HD-modulated (Fig. 7h,j), suggesting this coding may be behaviorally relevant. We therefore hypothesize that HD-modulated place cells may be more prevalent in the mammalian hippocampus moving in 2D than currently appreciated. Our simulations reveal that spatial representations are very sensitive to small changes in task requirement and constraints on behavior (cf. Vijayabaskaran and Cheng, 2022).

## 3 Discussion

We demonstrated that a computational model that flexibly learns to store and retrieve memories can solve auto- and hetero-associative memory tasks as well as a variety of spatial navigation tasks. The model spontaneously learns representations with task-related geometry in low-dimensional manifolds, e.g., categorical class representations in memory tasks similar to concept cells (Quiroga et al., 2005), attractor-dynamics-like representations of a morph sequence (Leutgeb et al., 2005; Wills et al., 2005; Colgin et al., 2010), highly specific representations of episodes, called barcodes (Chettih et al., 2024), ego-central goal representations (Sarel et al., 2017), and behavior-dependent head-direction modulation of spatial representation (McNaughton et al., 1983; Battaglia et al., 2004; Markus et al., 1995). In parallel, the model also learns complex computations over these representations and, in doing so, exploits the linear algebra computation inherent in activity propagation in feed-forward networks (*y* = Φ*x*). In particular, the model learned in the guidance task to use the memory retrieval process to compute the ego-centric goal position from the allocentric goal representation in the FWM memory and allocentric current position in the reading key, which includes a coordinate transformation.

It is useful to contrast how spatial representations emerge in the present model with how this occurs in other computational models that do not encode information in one shot. One class of models emphasizes the role of the hippocampus in spatial computation and presupposes some computational principle that ensures that different views at the same location are encoded similarly, but similar views at nearby locations are encoded differently, i.e., place- or grid-like responses. Principles that can achieve this are, for instance, path integration (Samsonovich and McNaughton, 1997; Banino et al., 2018; Cueva and Wei, 2018)) and the extraction of a slowly-varying latent variable from visual inputs during navigation (Franzius et al., 2007). Another class of models has shown that generic computational principles, such as reinforcement learning and backpropagation, are sufficient to generate place-cell-like responses (Vijayabaskaran and Cheng, 2022). What is new in our study is that we explicitly focus on the role of one-shot memory, namely the key–value memory system (FWM) that stores information from single experiences and is accessed flexibly at later time points. In our model, spatial representations emerge in the *memory structure* as a consequence of solving the navigation task, that is, the model discovers autonomously that it is useful to operate with and store spatial representations.

Three key features enable the model’s versatility: Firstly, flexible memory writing and reading mechanisms trained with gradient descent, which allows task-dependent memory usage. Secondly, the separation of key and value vectors in the memory matrix, which enables distinct encoding of different input features. Thirdly, the separation of timescales. FWM memory encodes information instantly with minimum interference, but stores it long term; the LSTM can blend information across the past few trials; and backpropagation adjusts network weights fairly slowly.

Despite its strengths, the model lacks direct anatomical constraints, complicating its biological mapping. Since the LSTM acts as a controller for the LTM, one option is to map them onto the prefrontal cortex (PFC) and the hippocampus, respectively, aligning with PFC’s role in gating hippocampal input (Navawongse and Eichenbaum, 2013). However, the hippocampal formation has rich recurrent connections, especially CA3, whereas the memory matrix is feedforward. Alternatively, due to its recurrent structure, the LSTM may correspond to CA3 and the memory matrix to CA1, since CA1 receives inputs from CA3 and is the primary output of the hippocampus (van Strien et al., 2009). Other model components, such as the delta learning rule or using separate keys for writing and reading, are biologically implausible and require further study. Furthermore, some well-known features are absent from the model, e.g. replay, which allows agents to learn offline from previous experiences (Zeng et al., 2023), and low-frequency theta oscillations, which are known to organize hippocampal spiking activity and coordinate memory encoding and retrieval (Buzsáki, 2002). Nevertheless, the emergence of hippocampus-like representations in the model suggests that it captures essential principles of hippocampal function, notwithstanding the exact mapping between the model and brain anatomy.

Another important limitation of the present model is the absence of an explicit temporal context signal. Although the network can generate sequences through learned pair-wise associations, it does not represent a continuously evolving temporal context of the type proposed in temporal context models of episodic memory (Howard and Kahana, 2002). Consequently, the model does not account for temporal contiguity effects or context-based reinstatement phenomena observed in human free-recall paradigms. Incorporating an explicit temporal context representation into the present framework is an important direction for future work.

Our modeling results suggest that the hippocampus is a flexible memory device, capable of representing task-relevant variables when behavioral performance demands it and learning is successful. This task-dependence could help explain why hippocampal representations are increasingly found to be modulated by various stimuli, e.g., fear associations (Moita et al., 2003), tone frequencies (Aronov et al., 2017; Zutshi et al., 2025), or the presence of conspecific (Omer et al., 2018). We predict that any variable that is task-relevant will eventually be represented by the hippocampus.

We conclude that episodic memory might be the primary function of the hippocampus and spatial representations are only one, albeit very important, type of representation that the hippocampus comes to exhibit because it is needed for solving a task.

## 4 Materials and Methods

The computational model consists of four main parts (Fig. 1a): A convolutional neural network (CNN), a long short-term memory (LSTM), a fast weight memory (FWM) matrix, and a downstream network (DS). The CNN pre-processes camera inputs; the LSTM learns to interact with the memory matrix by gradually adjusting its own weights over multiple exposures and, hence, acts as a memory controller; the FWM updates its weights after single exposures and acts as a long-term memory (LTM) store. The downstream network concatenates the outputs from the LSTM and the FWM matrix and converts them into the model output.

### 4.1 The Fast Weight Memory Model

At each timestep *t*, memory writing and reading occur consecutively. Memory writing consists of:

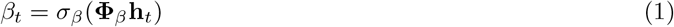

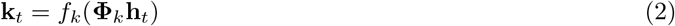

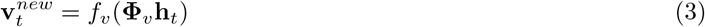

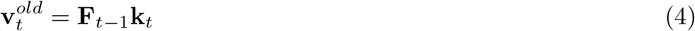

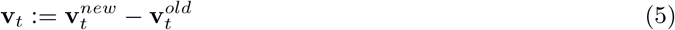

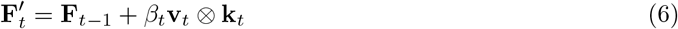

where **h**_*t*_ and **F**_*t*_ denote the LSTM output and the FWM matrix, respectively. *β*_*t*_ represents the writing strength, a scalar between 0 and 1, whose activation function *σ*_*β*_ is the sigmoid function. **k**_*t*_ and 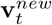 are the writing key and the new writing value vector, respectively. Throughout all equations, *f*_(·)_ refers to activation functions, and **Φ**_(·)_ refer to trainable weight matrices except for **Φ**_*β*_, which is a weight vector. We used ReLU activation functions for all key and value vectors to facilitate interpretation of a unit’s activation level as neural activity.

The update rule is termed a “delta learning” rule in the original paper (Schlag et al., 2021; Irie et al., 2021), because the outer product between the writing key and the “delta” writing value – the difference between the new and old writing value – is encoded into the memory and has been shown to improve memory capacity compared to simple addition. Throughout the paper, we refer to 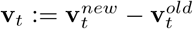 as the “writing value”. In addition, if we examine the update of the individual weights of the FWM matrix (eq. 6), 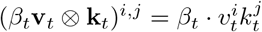 resembles the Hebbian learning rule, if the writing key and the writing value are considered the activity of a pre-synaptic and a post-synaptic layer.

Furthermore, to prevent the memory matrix from growing unbounded, the FWM matrix *F* is rescaled whenever its L2-norm *>* 2:

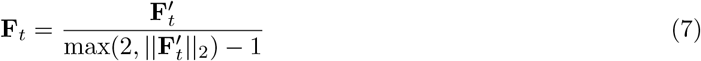

This is different from the original paper (Schlag et al., 2021) where the threshold is set to 1, leading to the denominator 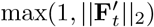. We found that this modification is crucial for the model to solve the two memory tasks, but not the spatial task. Although we did not perform any systematic analysis, we hypothesized that increasing the threshold for normalizing the memory matrix increased the memory capacity, which enabled the model to store the information of a sequence of images in the memory task. This was not so crucial in the spatial task where the model only needed to encode one piece of information: the goal location.

The reading process is governed by the following equations:

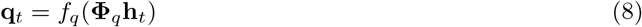

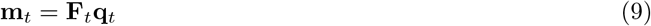

where **q**_*t*_ and **m**_*t*_ represent the reading key and value, respectively. The final output is computed by the downstream network (*DS*(·)) over the concatenation of the outputs from the LSTM and the FWM matrix:

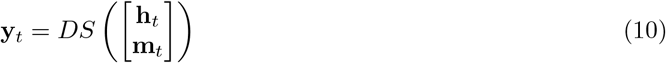

### 4.2 Model Hyper-Parameters

The CNN has three convolutional layers whose sizes depend on the tasks. In the memory task, the CNN receives gray-valued images of size 28 × 28 and consists of three convolutional layers (layer 1: 32 filters, 3 × 3, stride 1; layer 2: 64 filters, 3 × 3, stride 2; layer 3: 64 filters, 3 × 3, stride 2). In the navigation task, the CNN receives RGB images of size 84 × 84 × 3 and also consists of three layers, but with different sizes (layer 1: 32 filters 8 × 8, stride 8; layer 2: 16 filters, 4 × 4, stride 4; layer 3: 16 filters 2 × 2, stride 2). All layers use the ReLu activation function. The output of the CNN is flattened into a 1D vector and then used as the partial or complete input for the LSTM, depending on the task. The LSTM has a size of 256, and the FWM matrix has a dimension of 128 × 128 across all simulations.

In the spatial task, the LSTM also receives the representation of the previous action and the previous reward signal (Fig. 1f) as additional inputs, where the flattened output from the CNN (with a size of 256) is concatenated with an action vector and a reward vector. To make the different sources of inputs in the concatenation have comparable sizes, we encode the action by a one-hot vector (e.g. [1, 0, 0]) and duplicate the vector 100 times ([1, 0, 0, 1, 0, 0, …, 1, 0, 0]). Similarly, the reward is represented by a vector of size 100 whose elements share the same value, the immediate reward signal. In this way, the resulting input vector to the LSTM has a size of 256 + 300 + 100 = 656.

The downstream network (DS) in the memory task is a de-convolutional neural network with three transposed convolutional layers whose sizes match those in the CNN at the input level so that images have the same dimensions and the range of values as those of the inputs can be reconstructed. In the spatial task, the DS consists of two branches: an action branch and a value branch. The former is a linear layer with an output size 3, paired with the softmax activation function, to represent the distribution of selected actions. The value branch is a linear layer with output size 1, which represents the state value.

We used the RMSprop optimizer for the memory tasks and the Adam optimizer for the spatial tasks. The learning rate was 0.001 throughout the experiments.

### 4.3 Auto- and Hetero-Associative Memory Tasks

In the auto- and hetero-associative memory tasks, a trial refers to the presentation of one image to the model. The inputs to the model are randomly drawn from the EMNIST Dataset (Cohen et al., 2017) that includes 28 × 28 gray images of handwritten digits (0–9) and Roman letters (uppercase A – Z and lowercase a–z), in total 62 classes. To avoid class ambiguity, only the first 18 classes (digits 0-–9 and letters A–H) are used for training. The remaining 18 classes were used in the generalization study as retrieval cues. Both tasks are divided into an encoding phase and a retrieval phase, separated by a buffer trial between phases, in which a blank image was presented. The encoding phase is identical in both memory tasks: the model is presented a sequence of images of digits or uppercase letters. The retrieval phase differs between the tasks. Input sequence lengths vary between 8 and 18 across blocks. In the auto-associative memory task (Fig. 1c), the model has to output the exact encoded image based on a different cue image from the same class. In the hetero-associative memory task (Fig. 1d), original images are presented in random order, and the model has to output the next image that followed the cue in the encoded sequence.

During training (Fig. 1e), the entire sequence of the output images in the retrieval phase and the ground truth images are used to compute a binary cross-entropy loss (BCE) to update the model.

While reconstructing images from the writing and the reading value vectors, we replaced the LSTM output **h**_*t*_ with a zero vector in the stacked vector 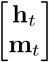 before passing it through the downstream network. To quantify unit responses to a given image class, we averaged the unit firing rates over all trials involving the images of that class in the test session.

### 4.4 Spatial Navigation Tasks

All spatial navigation tasks were simulated in the CoBeL-RL framework (Diekmann et al., 2023), a Python-based platform designed to flexibly simulate the navigation behavior of rodents by connecting powerful simulators, such as Blender and Unity with reinforcement learning (RL) algorithms. All maze environments were built in the Unity simulator. We adopted Proximal Policy Optimization (Schulman et al., 2017), a state-of-the-art, model-free RL algorithm known for stable training, to train the model on the spatial learning tasks.

#### 4.4.1 Maze-Morphing Experiment

In the maze-morphing experiment, the agent has to solve a navigation task in a series of mazes of different shapes (Supplementary Fig. 3). In the encoding phase, the agent has to search for an invisible reward. In the retrieval phase, the agent is rewarded for returning to the same location repeatedly until the end of the task block (Fig. 1f). This type of navigation behavior, in which agents rely on distal cues to locate themselves and the goal location is called “guidance” (Parra-Barrero et al., 2023). After reaching the goal location, the agent is transported to a random start location in the maze.

There are two training conditions: In the single-location condition, circle and square mazes are presented in the same location alternately in separate blocks; in the double-location condition, both mazes are present at different locations and connected by a corridor that allows traversal between them (Supplementary Fig. 3b). To make the circle and square mazes look the same in both conditions, the maze walls attached to the corridor in the double-location condition do not have any openings, but the agent can move through the wall into the corridor.

The agent receives a positive reward of +1.0 when reaching the goal and a small negative reward (− 0.01 in the square maze or 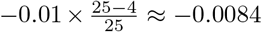 in the circle maze) at any node that is not the goal to penalize movements. There is, however, one exception. The agent receives 0 penalty when it reaches the four corners of the square maze, to model the preference of rodents to stay close to corners and to induce slightly different behaviors in the square and circle. To compensate for the four states that are not penalized in the square maze, the step penalties in the circle maze have been adapted to maintain the same average penalty across the two mazes. In the test session, the agent is tested on the guidance task in each of the six differently shaped mazes for 300 blocks.

##### Analysis of firing rate maps for individual units

The output of all units in the reading key are recorded while the agent naturally solves the task during retrieval, i.e., navigating to the goal node. For each of the units, its rate map across the maze is constructed separately for the four head directions (HD) as follows. At each node, the unit’s summed activity is divided by the total occupancy of the agent. The resulting rate map is divided by the maximal value across all units and blocks, so that each rate map is bounded between 0 and 1.

To evaluate how the units’ activity changes across the six mazes with the morphed shapes (Supplementary Fig. 3a), we computed the average rate across all locations of a maze and fitted the average rate as a function of the mazes with a sigmoid function (Colgin et al., 2010). Those that do not pass the fitting criterion (returning “nan” or unrealistically large values of variance) are filtered out, keeping those units whose average rates have a sharp transition along the morph sequence (Fig. 3c).

##### Population vector (PV) correlation

We constructed the population rate vectors by computing the mean of the reading key vectors at each node across all test sessions. While correlating the population vectors between two environments, we first correlated the PV’s for each corresponding pair of nodes and then computed the average correlation across all nodes.

#### 4.4.2 Barcode Experiment

In the experiment by Chettih et al. (2024), birds chose to store seeds at random sites when feeders were open and later retrieved them when feeders were closed. We model this experiment by alternating encoding and retrieval consecutively until a total of 2,000 steps are completed, which define a single episode. At each step, the agent can perform one of four actions: *move-forward, turn-left, turn-right*, or *open-site*. All actions incur a negative reward of -0.01, except when the agent opens the correct site, which gives a +1.0 reward and ends the encoding or retrieval phase. During encoding, the agent has to cache an item. Unlike the chickadee, our artificial agent has no intrinsic reason to cache it at a particular site, so we mark the cache site with a white sphere (Supplementary Fig. 6b). The agent therefore learns to navigate straight to the goal, using an aiming behavior (Parra-Barrero et al., 2023), and “cache” it with the open-site action. During retrieval, the agent is transported to a random starting position, and the cache site is visually indistinguishable from other locations, forcing the agent to rely on its memory of the cache’s location to retrieve the cache. The walls in the simulated environment are colored in four distinct colors matching the colors of the feeders used as cues in the original task, allowing the agent to orient itself relative to the walls using guidance (Parra-Barrero et al., 2023) (Supplementary Fig. 6a-b). In all five runs, the agents that were trained with the same hyper-parameters reliably learned to solve the task within ∼ 2,000 episodes (Supplementary Fig. 6c).

##### Population vector correlation

To reveal the barcode, we computed correlations between LSTM output vectors similar to how Chettih et al. (2024) analyzed the CA1 PV. Since the LSTM outputs project to both memory-writing (Eq. 6) and memory-reading pathways (Eq. 9), they provide a common representational space where cache, retrieval, and visit events can be directly compared. The mean PV for each event type is subtracted (Chettih et al., 2024), then the pairwise correlations between each cache event and all other cache, retrieval, or visit events are computed. For each pair, correlations are plotted as a function of the Euclidean distance between the agent’s positions at each event. Results were averaged over five trained agents, and variability was estimated via standard error of the mean (SEM).

##### Simplified synthetic model

We built a minimal synthetic model in which two 1D variables are jointly represented: the agent position (*a*) and goal position (*g*). Both variables are defined on a a 1D grid with five discrete locations (1–5). Events defined by (agent, goal) pairs are uniformly sampled 500 times (Fig. 4d, left). Events are labeled as *cache*, if *a* = *g* and *visit* otherwise. To make the events distinct, noise was added to each event *v* = [*a* + *ϵ*_*a*_, *g* + *ϵ*_*g*_], with *ϵ* ∼ 𝒩 (0, 0.2). We computed Euclidean similarity 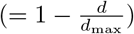, where *d* is the Euclidean distance, and averaged it within bins of agent-distance Δ*a* separately for cache-cache and cache-visit pairs (Fig. 4d, right).

##### Linear discriminant analysis (LDA) and ablation analysis

We applied two separate two-component Linear Discriminant Analyses (LDA) to the *d*-dimensional LSTM output: one using agent positions as class labels and one using goal positions as class labels. Each LDA learns a linear projection that maximizes between-class separation and provides a *d* × 2 weight matrix, where row *i* contains the two weights assigned to LSTM unit *i* across the two discriminant axes (the unit’s loading vector). To quantify how strongly unit *i* contributes to discrimination, we take the Euclidean norm of that two-element row vector. The “LDA norm” is computed separately for the agent- and goal-based LDAs (Fig. 4e).

To measure redundancy, we ranked units by their LDA-norms and removed either the *k* top-ranked units or *k* randomly selected units, where *k* = 0, 10, …, *d*. We then retrained LDA and measured classification accuracy for the agent and goal decoding tasks. This process was repeated 20 times and the results were averaged (Supplementary Fig. 6 d).

#### 4.4.3 Generalization in the Guidance Task

To test the generalization ability of the memory model in the simple guidance task, we removed two goal locations, node 4 and 17, from the training session. During the test session, all nodes in the maze were used as goal location. We compared the latency of finding the novel goal nodes to that of familiar goal nodes that are equally reachable under a random policy. For instance, it is less probable to reach a corner node than the central node. Hence, goal node 4 is compared to goal node 6, 20 and 22; and goal node 17 is compared to goal node 7, 9 and 19 (see Supplementary Fig. 10a for the numbering).

#### 4.4.4 Continuous State Space with Discrete Tank Actions

In addition to the grid-based navigation tasks, we implemented a continuous-state spatial navigation task based on a tank robot model. The agent moves in a continuous 2D space (5 × 5 unit square), but controls its motion through three simple discrete actions: (1) turn right (left wheel active, right wheel stopped), (2) turn left (right wheel active, left wheel stopped), (3) move forward (both wheels active). Each action updates the agent’s pose (*x, y, θ*): forward steps translate the agent by 0.2 units along its heading direction, while turning actions rotate the agent by ≈ 23° and produce the corresponding arc-shaped movement characteristic of the tank model. The agent is considered to have arrived at the goal location, if it is positioned within a distance of 0.8 units from the goal.

#### 4.4.5 Classification into Place or Head-Direction Modulated Units

To categorize the selectivity of the units in both the discrete and continuous state space experiments (Fig. 7), spatial rate maps are computed separately for different agent orientations. In the discrete state space, four HDs are possible and for each a 5 × 5 firing rate map is computed. In the continuous state space, HD is binned into eight 45° intervals centered at 0°, 45°, …, 315° and, for each HD, a firing rate map is extracted for 10 × 10 bins. Each map reflects the average activity of the unit at each spatial location while the agent is facing the corresponding direction.

Two metrics extracted are used to access spatial representations: Unrotated spatial similarity (*S*_space_), which is the mean pairwise correlation across all directional maps, and rotation-aligned similarity (*S*_rot_), which is computed similarly to the previous measure, except the maps are first rotated by their orientation to align them. Units with *S*_space_ *> S*_rot_ and *S*_space_ ≥ 0.15 (4-action) or ≥ 0.25 (continuous) are labeled *place units*; those with *S*_rot_ *> S*_space_ and *S*_rot_ ≥ 0.15 (4-action) or ≥ 0.25 (continuous) are labeled *conjunctive units*; units that do not meet these criteria, or have peak firing *F*_max_ < 0.25 in all maps, are labeled as *non-spatial*.

## 5 Acknowledgements

Funded by the Deutsche Forschungsgemeinschaft (DFG, German Research Foundation), project number 397530566 (FOR 2812, P2 and P5).

## 6 Supplementary Text

### 6.1 Emergent Geometric Computation in Spatial Learning Task

Spatial computations emerge in our model through the combination of how the long-term memory (LTM) matrix is constructed in the fast weight memory (FWM) model and the geometry of the representations that the model learns autonomously. Here, we analyze the first retrieval phase in a task block (between the first and second time that the agent reaches the goal), but the results generalize to all other trials. We will show below how the egocentric representation of the goal location in the reading value **m**_*t*_ in the retrieval phase (Fig. 5i) emerges from the interaction between the representation of the goal location in **k**_*g*_ when the agent first reaches the goal (Fig. 5c) and the representation of the agent’s location in **q**_*t*_ (*t* > *g*) during retrieval (Fig. 5g,h). We break down this explanation into four steps:

1. The direction and magnitude of the reading value **m**_*t*_ during retrieval is dominated by the scalar product between the writing keys (**k**_*g*_, **k**_*g*+1_) when the agent is at the goal and one timestep later, respectively, and the reading keys **q**_*t*_ during retrieval.
2. The geometry of the representations of **k**_*g*_ and **q**_*t*_ are such that their scalar product in PCA space indicates the egocentric goal location along the sagittal axis (front-back).
3. The ReLU activation in the network makes all key/value vectors non-negative. Hence, the sign of **k**_*g*_ · **q**_*t*_, computed on actual vectors, is non-negative and cannot be used to indicate relative locations. The model has to solve this problem.
4. The second PCA component of **v**_*g*+1_ encodes the goal position along the transverse direction.

#### Step 1

Equations 6 and 7 allow us to rewrite the reading output at step *t* as follows:

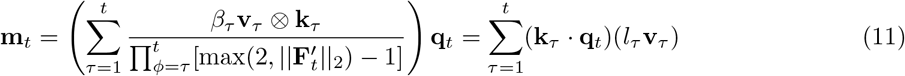

where 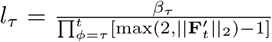 is a scalar. Plotting *l*_*τ*_ from *τ* = 0 to *T*, where *T* is the last step of the first retrieval phase, shows that *l*_*τ*_ is negligible in the encoding phase except for at step *g* (Supplementary Fig. 7a). This implies that the information encoded into the memory during exploration does not contribute to the reading output during retrieval. Intuitively, this is the consequence of the combination of medium-sized beta values and the frequent normalization of the FWM matrix (Eq. 7). Additionally, *l*_*τ*_ is also negligible during the retrieval phase other than at step *g* + 1. This allows us to simplify the equation to:

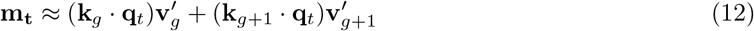

where 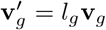 and 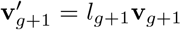. To validate our analysis, we performed PCA on **m**_**t**_ constructed according to the simplified Eq. 12 and found the same structure as in the network output (Supplementary Fig. 7i, left).

#### Step 2

To see how Eq. 12 helps in understanding the geometric computation in the network, we projected writing keys **k**_*g*_ and reading keys **q**_*t*_ in one PCA space. We found that they are approximately located on the same sphere (Supplementary Fig. 7b). In the 2D PCA space, the clusters representing the Y coordinate of the agent’s position while heading north and south (reading key) are approximately located on the same circle with those clusters representing goal locations (writing key) (Supplementary Fig. 7c). A further investigation revealed that there is a ±90° offset between this pair of clusters (sketched in Supplementary Fig. 7d for clarity). This geometric structure of the key representations turns out to enable the computation of the relative goal location along the sagittal axis (front-back) of the agent. For example, let us take the case where *y*_goal_ = 5 and *y*_agent_ = 3 (Supplementary Fig. 7d, e). If the HD is north, the configuration results in a positive scalar product between the writing and reading keys, **k**_*g*_ · **q**_*t*_ *>* 0, i.e., *α* < 90^*o*^ (Supplementary Fig. 7d, solid blue line and solid green line). If, on the other hand, the agent is heading south, the scalar product is negative (Supplementary Fig. 7d, dotted blue line and solid green line). Hence, the sign of the scalar product between the PCA projections of **k**_*g*_ and **q**_*t*_ indicates whether the goal is in the front or back of the agent, and the magnitude of the scalar product indicates the distance to the goal. The average values of the above scalar products for different *y*_goal_ while fixing *y*_agent_ = 3 indeed show this pattern (Supplementary Fig. 7f, solid orange line).

#### Step 3

There is however a small issue with this elegant geometric computation: The computations in the model are performed in the neural network, not in PCA space. In the neural networks, the key vectors are all non-negative as a result of the ReLU activation function. Hence, the scalar product between key vectors can never be negative (Supplementary Fig. 7f, blue curves). The model develops an interesting solution to this problem. First, since there are two scalar products in Eq. 12, their summation could be used to subtract the right contribution to correct for the non-negativity of vectors. For this to work, the two vectors in Eq. 12, 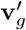 and 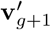, must be aligned in opposite directions. However, since the vectors are high-dimensional it suffices if one direction is aligned, e.g., the first principle component (PC). Let 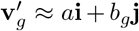 and 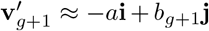, where **i, j** refer to the first and the second PC direction, respectively, the reading output **m**_*t*_ can be rewritten as follows:

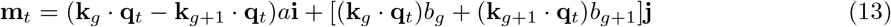

PCA analysis on the normalized value vectors 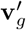 and 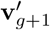 show that indeed their first PCs point in opposite directions (Supplementary Fig. 7g, h). Then, the curve of the scalar product based on the key vectors and that based on the PCA projections are almost parallel to each other (Supplementary Fig. 7f, blue and solid orange curves), meaning a constant value should be subtracted from the former to get the latter. Interestingly, the scalar product between **k**_*g*+1_ · **q**_*t*_ turns out to be constant (Supplementary Fig. 7f, black curves). At last, subtracting the two scalar products *p* = **k**_*g*_ · **q**_*t*_ − **k**_*g*+1_ · **q**_*t*_ restores the signed representation (Supplementary Fig. 7f, dashed orange curves). Indeed, the sign of *p* reliably predicts whether the agent moves forward or turns (Supplementary Fig. 7i, right).

#### Step 4

Finally, the second term in Eq. 13 that corresponds to the second PC is used to encode the relative goal location along the transverse axis and compute the correct turn. Specifically, the projections of the writing values at step *g* + 1 (the first step in the retrieval phase) along the second PC, *b*_*g*+1_, already reflects how far the goal lies to the right/left side of the agent (Supplementary Fig. 7j). To formalize this according to equation 13, we let *C*_1_ = (**k**_*g*_ · **q**_*t*_)*b*_*g*_ and *C*_2_ = (**k**_*g*+1_ · **q**_*t*_)*b*_*g*+1_. *C*_2_ naturally carries the same information as *b*_*g*+1_ because **k**_*g*+1_ · **q**_*t*_ is close to a constant (Supplementary Fig. 7f, black curve). On the other hand, *C*_1_ + *C*_2_ also carries similar information (Supplementary Fig. 7j), since *C*_1_ has relatively low magnitude compared to *C*_2_ and therefore does not substantially alter the representation.

In this way, we show that the relative goal location along the transverse axis is already encoded in the writing value **v**_*g*+1_, which is essentially generated by projecting the LSTM output through a weight matrix. Hence, we suppose the actual computation of this information is carried out by the LSTM.

### 6.2 Functional Role of Spatial Representations and Computation

Having explained how spatial computations are performed using the fast weight *memory* network, we next ask whether the emergent spatial representations and computations are functional or just epiphenomena. We therefore examined two conditions. First, during learning the pattern of the scalar products between keys developed gradually during the training session. This development is accompanied by the emergence of spatial representations in the PCA space and an increased performance of the agent on the task (Supplementary Fig. 8). The parallel emergence of the scalar products, representational structure, and task performance strongly suggest that there is a causal link between these variables. Second, the agent was able to generalize the spatial computations to held-out goal locations. During the test session, the agent could find and go back to goal nodes that it had never been trained on (Supplementary Fig. 10c). This generalization, even though performance was slightly lower, was realized through the geometric computation, because the representations of the novel goal nodes in the memory matrix appeared at the consistent locations in the representation space (Supplementary Fig. 10b).

To summarize, this theory demonstrates how a memory network and the geometric structure of memory representations supports spatial computation. These spatial representations and computations are not mere correlations, but have a causal role in the goal-directed behavior of the agent.

## 7 Supplementary Figures

**Figure 1:**
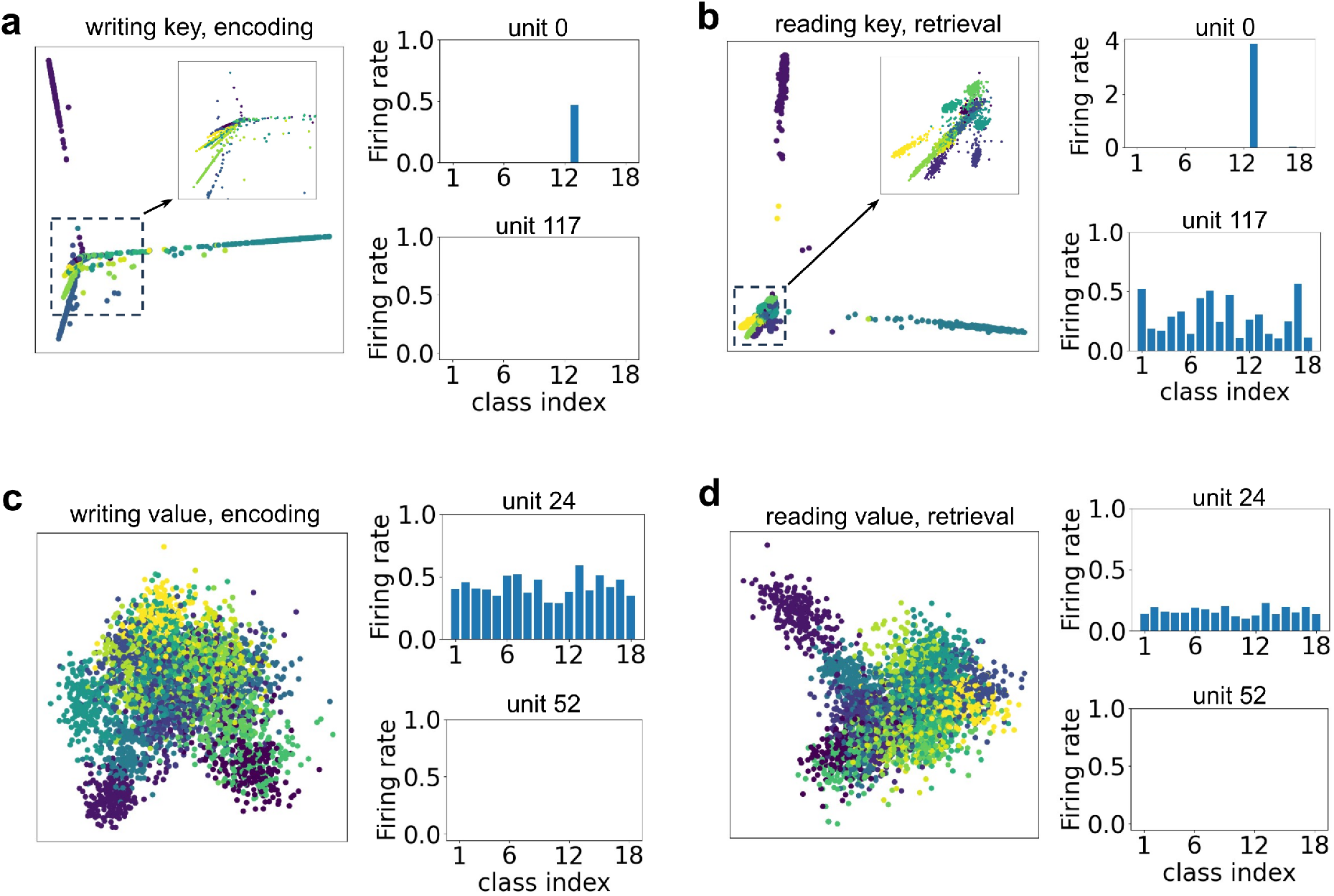
Mnemonic representations in auto-associative memory task. **a.** PCA projections of the writing key in the encoding phase (left) and examples of single-unit activation, a.k.a. the firing rate, for two units (right) when images of different classes are presented to the model. All PCA projections are colored w.r.t. the 18 classes. The units in the writing key are either categorical (unit 0) or silent (unit 117). **b**. Same as in panel a for the reading key in the retrieval phase. Units in the reading key reacts either categorically to the same class (unit 0) as the corresponding unit in the writing key or near-uniformly to all classes (unit 117). In this way, the dot product between a writing key and a reading key vector will only be non-zero, if their represented image classes match. **c**. Same as in panel a for writing value. **d**. Same as in panel a for reading value.

**Figure 2:**
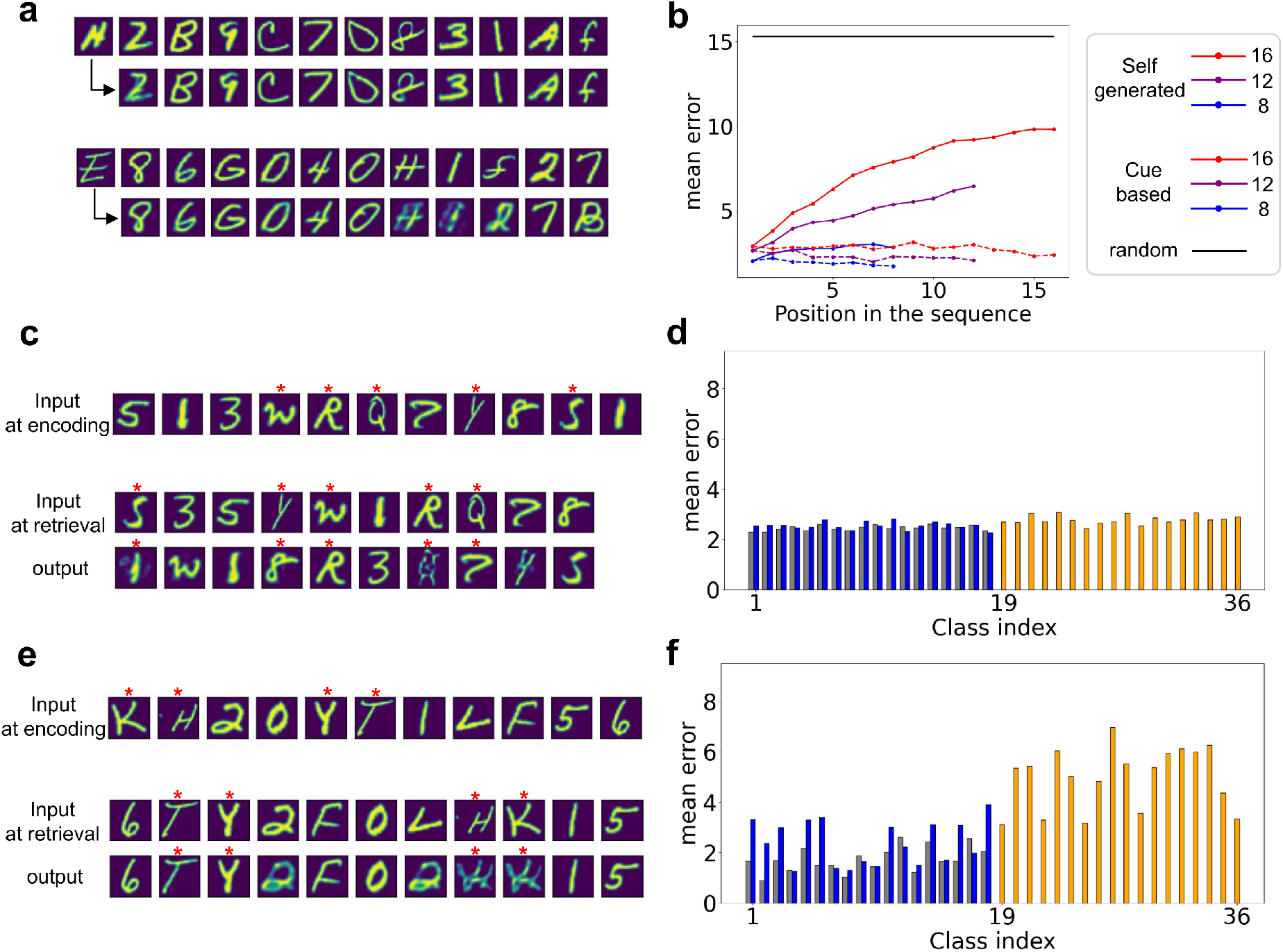
Two types of generalization in memory task. **a.** Two examples of self-generated sequences by the model in the hetero-association memory task, where the first row is the input sequence during encoding and the second row is the generated sequence, when the first image was presented as a retrieval cue. **b**. Mean errors between the generated image and the ground truth as a function of the position of the image in the sequence. **c**. An example test block of the generalization experiment for the hetero-associative memory task, where images whose classes were never presented in a training session were used in the testing session (red stars). The model can encode and retrieve images that were never seen before. **d**. The average error between the output image and the ground truth w.r.t. the class index for the hetero-association memory task. The first 18 classes (blue) were used in the training, while the last 18 classes (orange bars) were not. The gray bars represent the control situation, where no unseen classes were presented. **e, f**. The same figures as panels c, d, but for the auto-association memory task.

**Figure 3:**
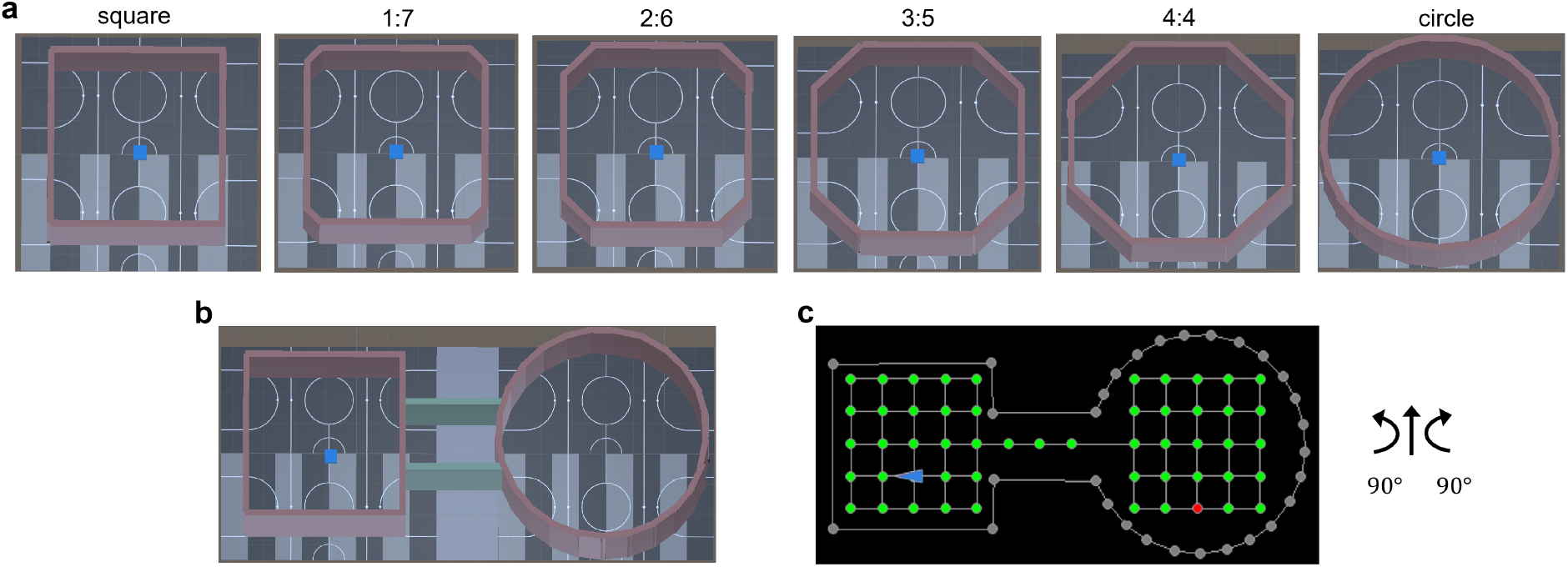
Simulated environments built in Unity for the maze-morphing experiment (Colgin et al., 2010). **a**. From left to right: Square maze, Maze 1:7, Maze 2:6, Maze 3:5, Maze 4:4, Circle maze. In each panel, the blue cube represents the artificial agent. The ratios (1:7, 2:6, etc.) represent the ratio between the length of the short side walls and that of the long side walls. **b**. The maze used in the training session of the double-location condition. **c**. Left: The topology graph in the double-location condition. The green nodes represent potential starting locations and the red node marks the current goal node; the blue arrowhead indicates the location and head direction of the agent. Right: Sketch of the action space. At each timestep, the agent can move forward, or turn 90° to the left or right. The agent remains on the same node if the chosen action would take it outside the maze.

**Figure 4:**
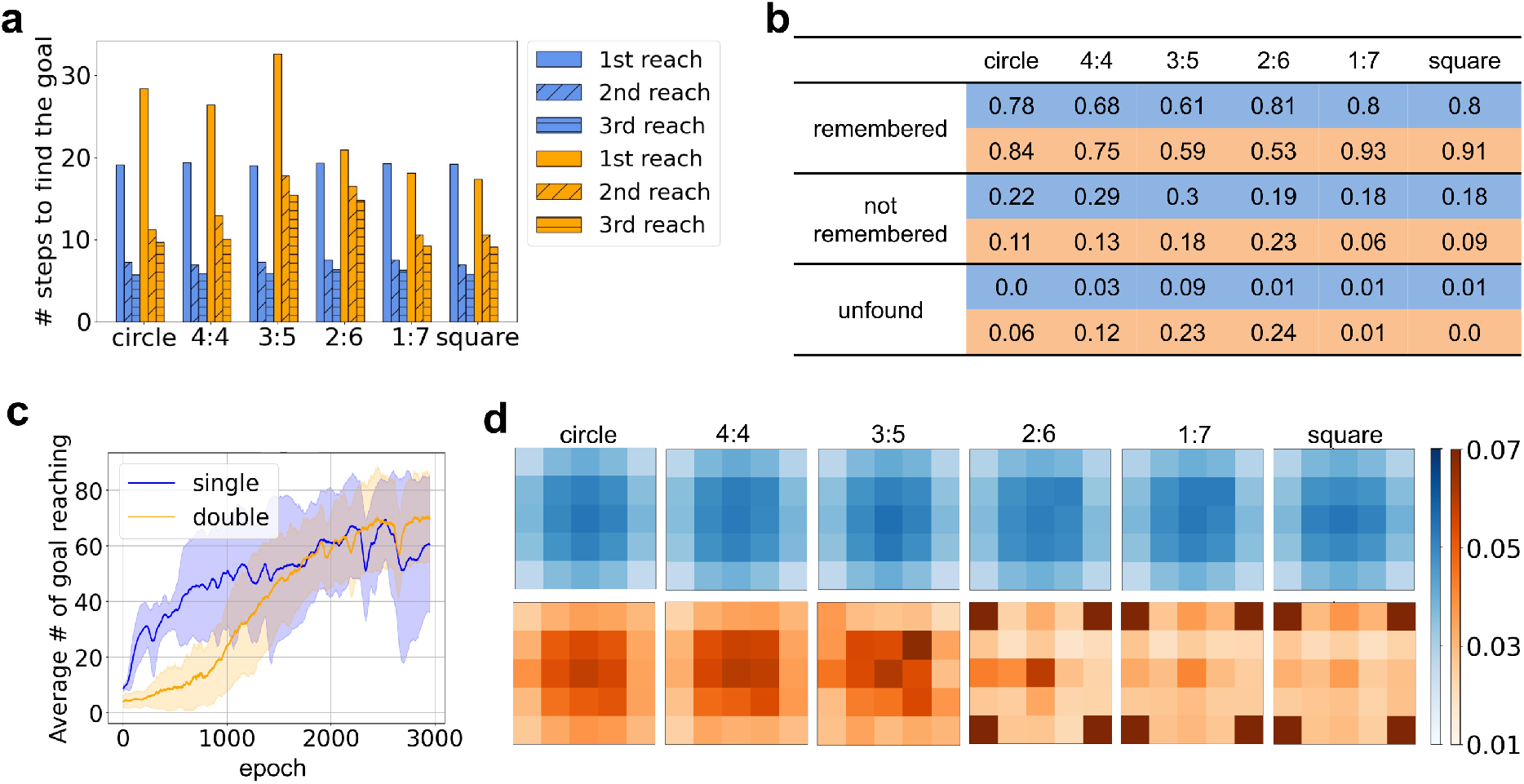
Behavioral analysis in the maze-morphing experiment. In all panels, blue color indicates the single-location training condition while orange color represents the double-location training condition. **a**. Average number of steps for the agent to reach the goal in the first three trials. The agent immediately took fewer steps after reaching the goal for the first time, indicating one-shot learning of the goal location. **b**. Numbers in the table represent the proportion of retrieval blocks where the goal location is remembered, not remembered, or unfound. **c**. Learning curves of the agent during the training session. **d**. Average rate of node occupancy by the agent in each maze in the two conditions. The agent showed a relatively homogeneous exploration patterns in all mazes in the single-location condition. By contrast, in the double-location condition there is a clear preference for the four corners in mazes 2:6 and 1:7, and in the square maze.

**Figure 5:**
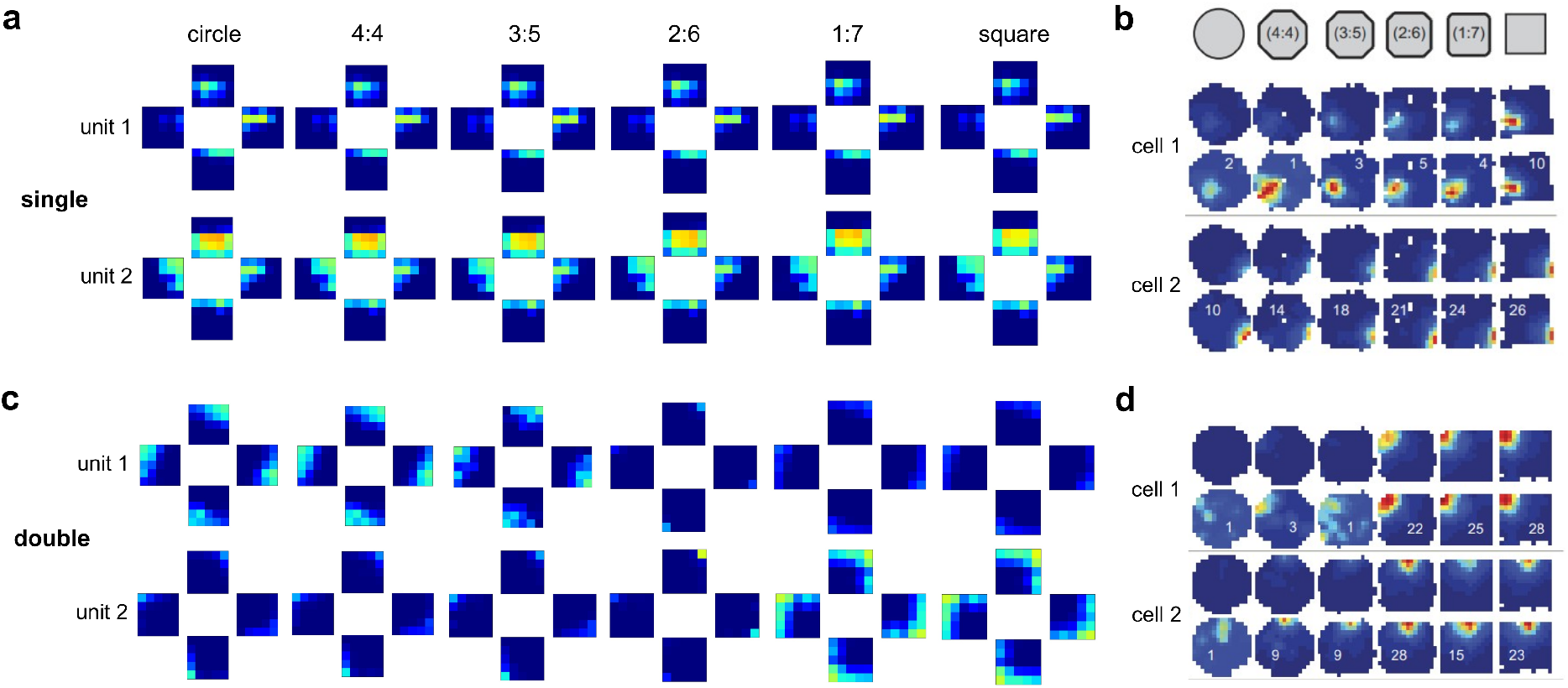
Single-unit activity maps in the maze-morphing experiment. **a.** Two example units in the reading key that show spatial tuning during the single-location training condition. For each combination of unit and maze, the four rate maps represent the response of the unit when the head direction of the agent is north, east, west, and south. The rates of all units have been normalized by dividing by the maximum firing rate across all sessions. **b**. Two example CA3 cells from Colgin et al. (2010) in the single-location condition, reproduced with permission. For each cell, the upper row depicts the firing rate in its absolute value, while the bottom row shows it after normalization, with the number representing the peak firing rate. **c, d**. The same as panels a, b, but for the double-location training condition.

**Figure 6:**
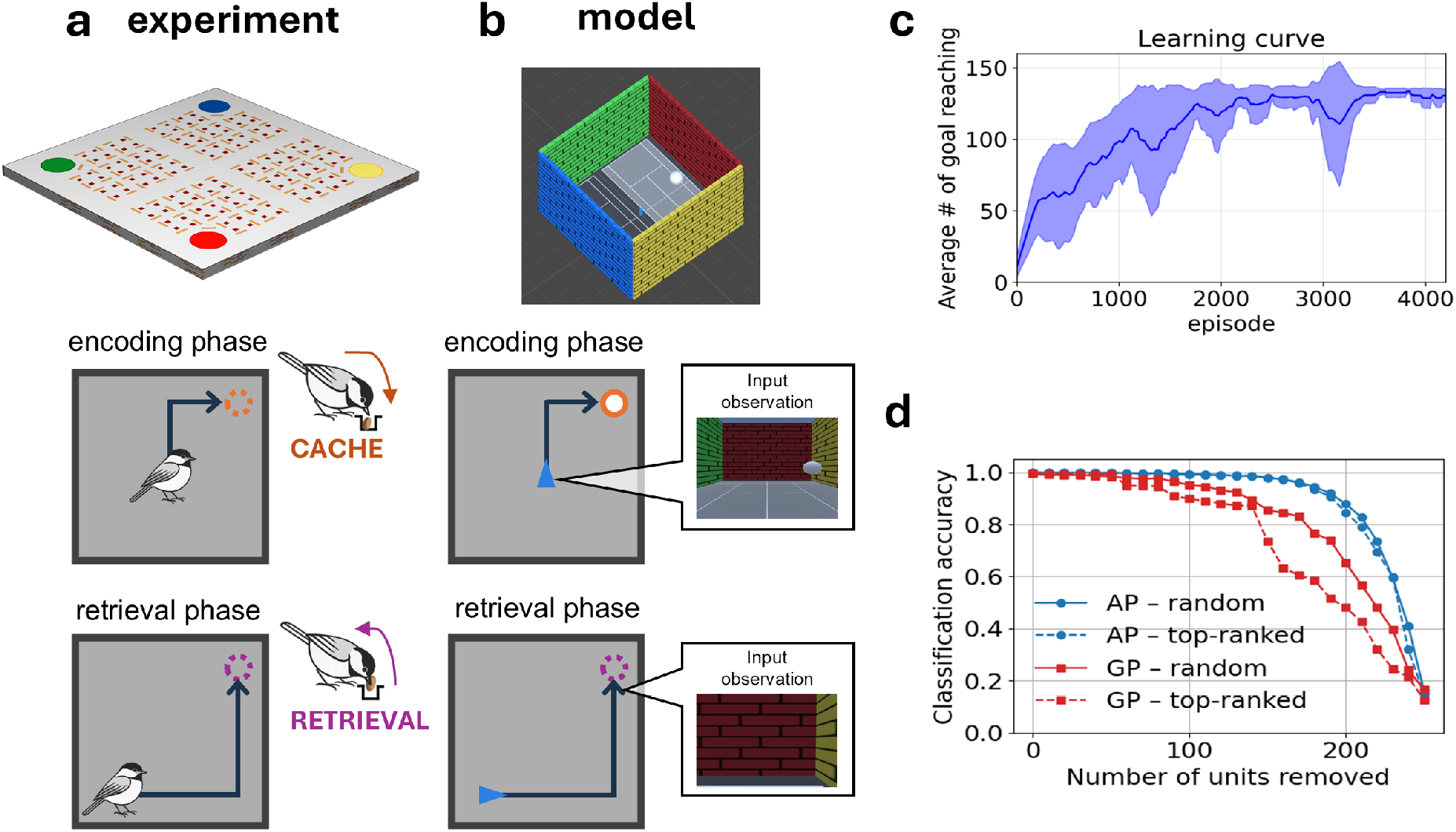
Setup of barcode experiment and simulation (Chettih et al., 2024). **a.** Schematic of behavioral task. Top: Overhead view of the maze containing four colored feeders. Bottom: simplified illustration of the trial structure consisting of an encoding and retrieval phase. **b**. Illustration of the corresponding simulation in our study. Top: 3D rendering of the square maze with four colored walls matching the feeder cues. Bottom: illustration of the trial structure with the agent’s egocentric perspective that is given as an input to the model. The agent moves through the environment using three spatial actions (*move forward, turn left*, and *turn right*). An additional action *open-site* is analogous to the chickadee lifting the cover of a cache site and serves as both a caching and retrieval mechanism depending on the task phase. Corresponding to the feeders-open period in the experiment, during encoding, the agent has to navigate to a visible goal, a white sphere, and “cache” food with the *open-site* action. Similar to the feeders-closed period in the experiment, during retrieval, the agent starts from a random location and must return to the cache site while the goal is not visible. After arriving at the cache site, the agent has to perform the *open-site* action to receive a reward. Once the cache is retrieved, the block ends and the next block starts with a new cache location (Fig. 1g). **c**. Average learning curve averaged across five independent agents. Shaded region represents the standard deviation. **d**. Classification accuracy of LDA decoding for agent-position (AP, blue) and goal-position (GP, red) as a function of the number of LSTM output units removed. The *k* deleted units are chosen either randomly (solid lines) or from the units with highest LDA loading, top-ranked (dashed lines).

**Figure 7:**
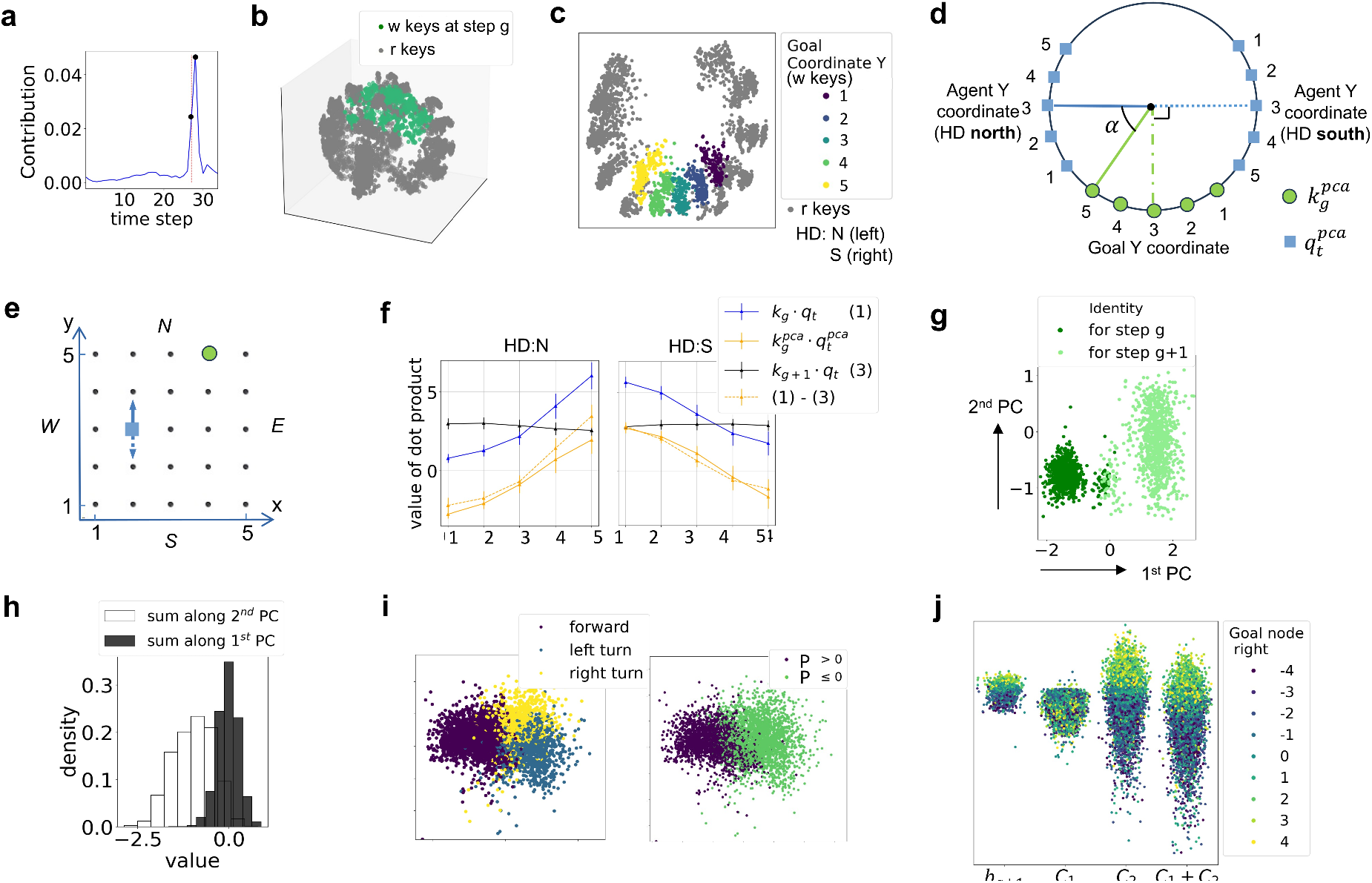
Demonstration of the spatial computation based on the geometric structure of the memory representations in navigation task. **a.** The value of *l*_*τ*_ from Eq. 11 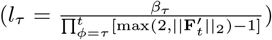 in an example test block. Here *t* in the formula of *l*_*τ*_ is taken to be the last step of the first retrieval period (the last point of the x axis). Step *t* = *g* and *t* = *g* + 1 are marked with black dots on the curve. **b**. 3D PCA projections of writing keys **k**_*g*_ and reading keys **q**_*t*_ during the retrieval phase are approximately located on the same sphere. **c**. 2D PCA projections of the same components as in panel b, where only the reading keys corresponding to head direction north (HD=N) and south (HD=S) are used for the projections. The projection of the writing keys has been color-coded based on the Y coordinates of the goal locations. The color code of the reading key has been demonstrated in Fig. 5h and is not shown here for clarity. **d**. A schematic for the observed geometric structure in panel b, depicted for the configuration in panel e. Blue squares represent the projection of reading keys that encode the Y coordinate and the HD of the agent, while the green circles represent the writing key projections that encode the Y coordinate of the goal location. **e**. An example configuration of goal location, agent location, and HD. Since the agent moves forward when the goal is in front of it, and turns left/right when the goal is behind it, the agent will move forward at this moment if it faces north, and will turn left if it faces south. **f**. Values of scalar products between the writing keys **k**_*g*_ and **k**_*g*+1_, and those of the reading keys **q**_*t*_ for *y*_agent_ = 2, averaged over the X coordinates of both the agent and the goal. **g**. First principle component (PC) of normalized writing values **v**_*g*_ and **v**_*g*+1_ point in opposite directions. **h**. Sum of the coordinates of the two writing values along the first PC are centered around 0, meaning that they have opposite values. **i**. Left: PCA projections of vectors computed from Eq. 12, colored by the action taken by the agent. Right: Same projections as on the left, but colored based on the sign of the value of *p* = **k**_*g*_ · **q**_*t*_ − **k**_*g*+1_ · **q**_*t*_ (cf. Eq. 13). **j**. Values of *b*_*g*+1_, *C*_2_ and *C*_1_ + *C*_2_ from Eq. 13 all contain information of the relative goal location in the transverse axis. Positive values in the color legend mean the goal is on the right of the agent, while negative values mean the goal is on the left.

**Figure 8:**
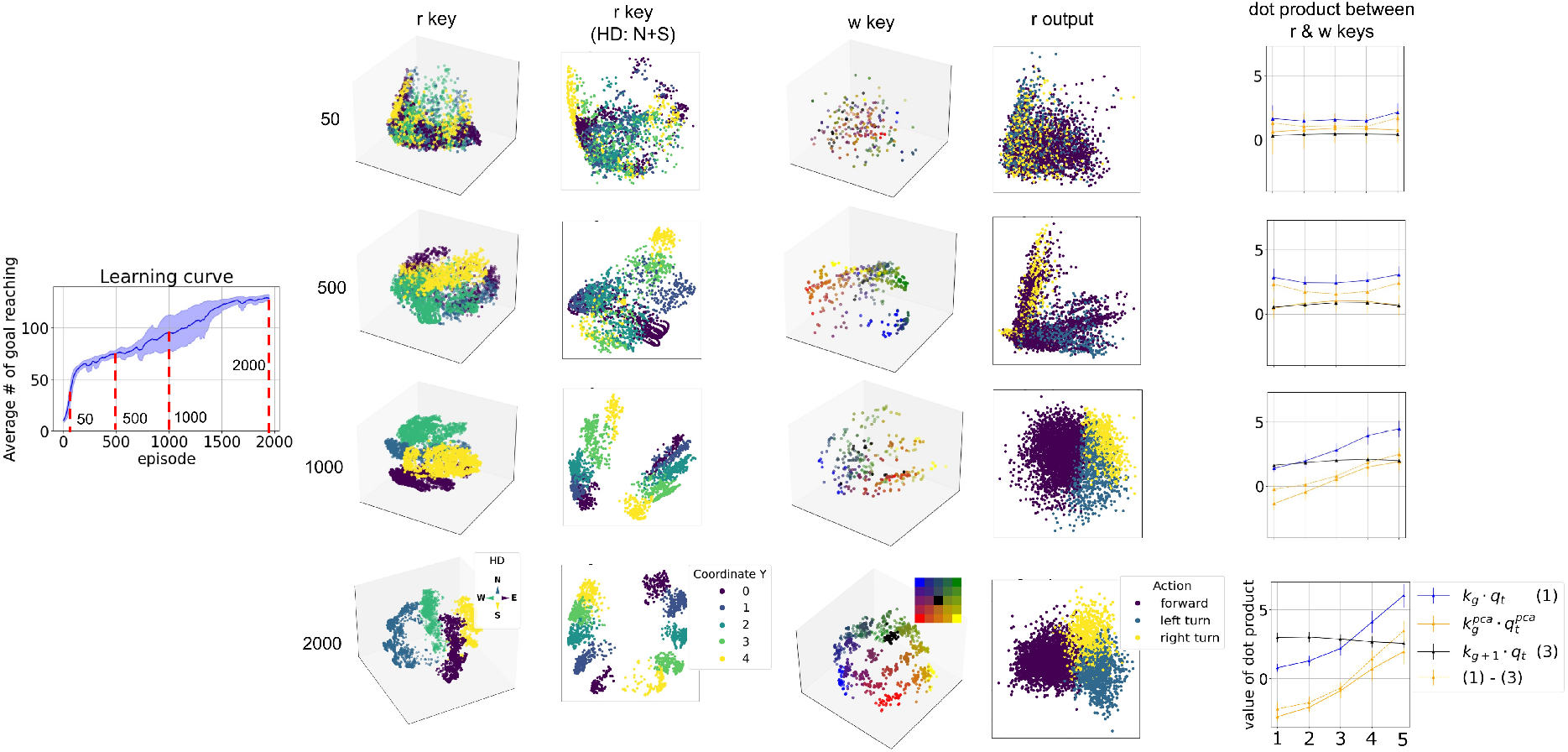
Emergence of geometric structure and performance improvement in the guidance task during training. First column: Learning curve. The model was tested at different training stages at 50, 500, 1000 and 2000 episodes (rows). Second-fifth column: Evolution of the spatial representations in the reading/writing keys and the reading outputs. Plotting conventions are as in Fig. 5g, h, c, and i. Last column: Evolution of the scalar products between writing keys and reading keys as in Supplementary Fig. 7f.

**Figure 9:**
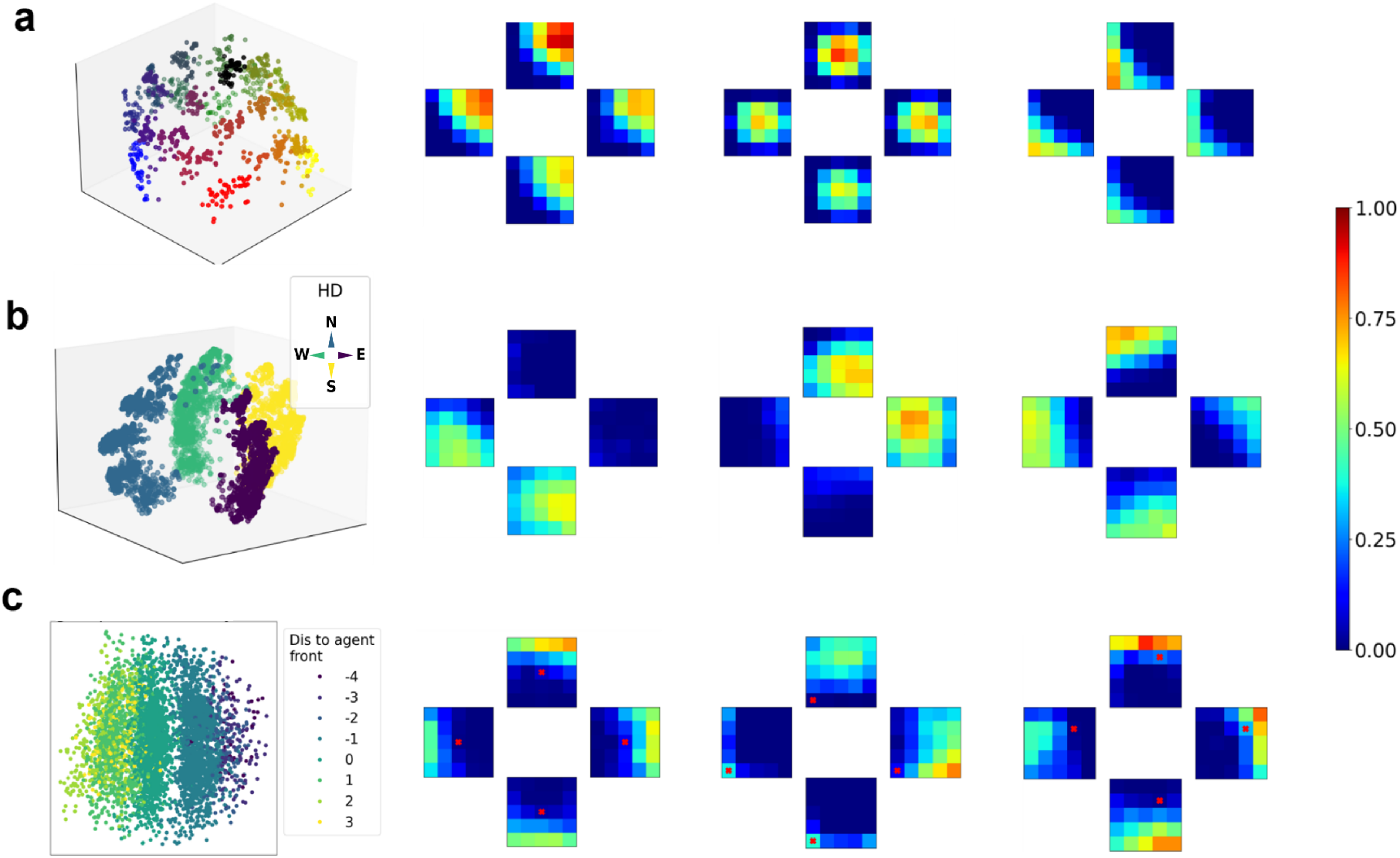
Examples of single unit activity maps in the guidance task in the square maze. **a.** PCA projection and three example units of the writing key when the agent is at the goal location. Data points in projection are colored by goal location as in Fig. 5c. Activity maps are plotted using the same convention as in Supplementary Fig. 5. Hence, each cluster in the projection and each small square in the activity map corresponds to a particular goal location. **b**. PCA projection as in Fig. 5g and three example units of the reading key in the retrieval phase. Each small square in the activity map corresponds to a particular agent location **c**. PCA projection and three example units of the reading output (value) in the retrieval phase. In the projections, the coloring corresponds to the relative position of the goal to the agent along the head direction. For each activity map, the goal location (red cross) is fixed while the activity is recorded, and each small square corresponds to an agent location.

**Figure 10:**
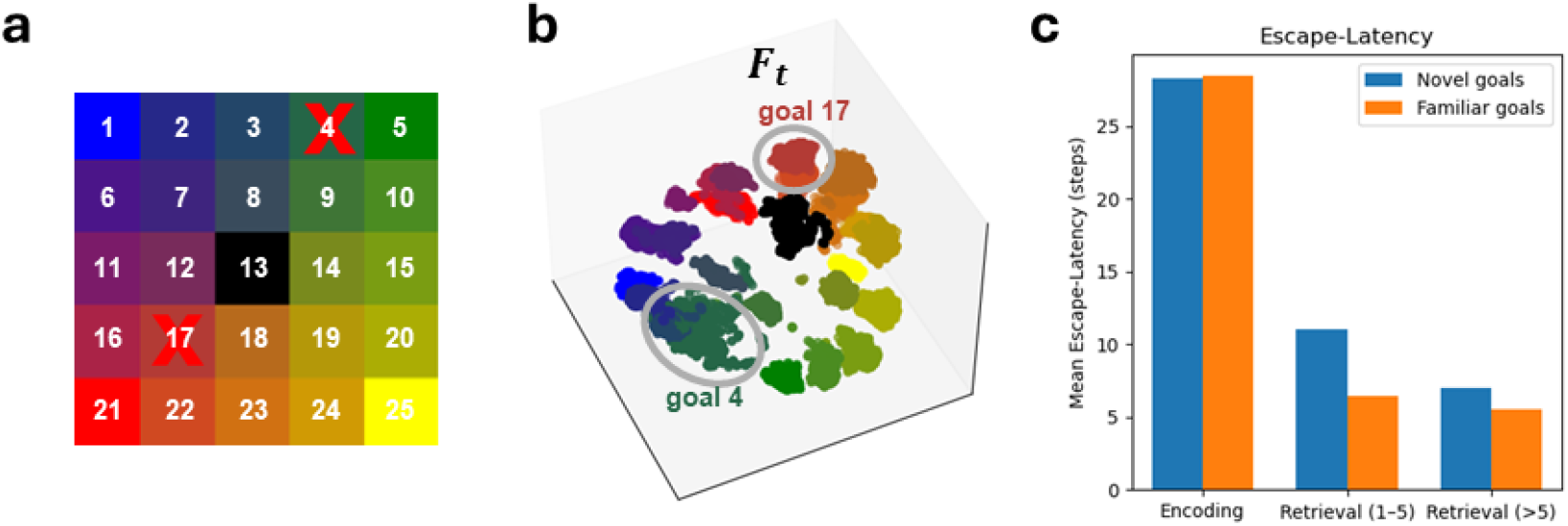
Generalization to novel goals in the guidance task. **a.** Numbering of the nodes. Nodes with red X were not used for training in the generalization task. **b**. PCA representations of the FWM matrix **F**_*t*_ showing that novel goals (circled: nodes 4 and 17) fall into the same manifold as nearby goals, forming distinct clusters even though the locations were never used as goals during training. **c**. Mean escape latency across different phases: during encoding, the agent takes a similar number of steps to reach both novel and familiar goals; during the first five retrieval attempts, latency is slightly higher for novel goals; later the performance on novel goals converges to that for familiar goals.

## References

Aronov, D., Nevers, R., and Tank, D. W. (2017). Mapping of a non-spatial dimension by the hippocampal– entorhinal circuit. Nature, 543(7647):719–722.

Banino, A., Barry, C., Uria, B., Blundell, C., Lillicrap, T., Mirowski, P., Pritzel, A., Chadwick, M. J., Degris, T., Modayil, J., Wayne, G., Soyer, H., Viola, F., Zhang, B., Goroshin, R., Rabinowitz, N., Pascanu, R., Beattie, C., Petersen, S., Sadik, A., Gaffney, S., King, H., Kavukcuoglu, K., Hassabis, D., Hadsell, R., and Kumaran, D. (2018). Vector-based navigation using grid-like representations in artificial agents. Nature, page 1.

Battaglia, F. P., Sutherland, G. R., and McNaughton, B. L. (2004). Local Sensory Cues and Place Cell Directionality: Additional Evidence of Prospective Coding in the Hippocampus. Journal of Neuroscience, 24(19):4541–4550.

Buzsáki, G. (2002). Theta oscillations in the hippocampus. Neuron, 33(3):325–340.

Chandra, S., Sharma, S., Chaudhuri, R., and Fiete, I. (2025). Episodic and associative memory from spatial scaffolds in the hippocampus. Nature, pages 1–13.

Cheng, S. (2013). The CRISP theory of hippocampal function in episodic memory. Front Neural Circuits, 7(May):88.

Chettih, S. N., Mackevicius, E. L., Hale, S., and Aronov, D. (2024). Barcoding of episodic memories in the hippocampus of a food-caching bird. Cell, 187(8):1922–1935.e20.

Cheu, E. Y., Yu, J., Tan, C. H., and Tang, H. (2012). Synaptic conditions for auto-associative memory storage and pattern completion in Jensen et al.’s model of hippocampal area CA3. Journal of Computational Neuroscience, pages 1–13.

Cohen, G., Afshar, S., Tapson, J., and van Schaik, A. (2017). EMNIST: An extension of MNIST to handwritten letters. 1702.05373.

Colgin, L. L., Leutgeb, S., Jezek, K., Leutgeb, J. K., Moser, E. I., McNaughton, B. L., and Moser, M.-B. (2010). Attractor-Map Versus Autoassociation Based Attractor Dynamics in the Hippocampal Network. Journal of Neurophysiology, 104(1):35–50.

Cueva, C. J. and Wei, X.-X. (2018). Emergence of grid-like representations by training recurrent neural networks to perform spatial localization. arXiv, page 1803.07770.

de Camargo, R. Y., Recio, R. S., and Reyes, M. B. (2018). Heteroassociative storage of hippocampal pattern sequences in the CA3 subregion. PeerJ, 6:e4203.

Diekmann, N., Vijayabaskaran, S., Zeng, X., Kappel, D., Menezes, M. C., and Cheng, S. (2023). CoBeL-RL: A neuroscience-oriented simulation framework for complex behavior and learning. Frontiers in Neuroinformatics, 17:1134405.

Eichenbaum, H. (2017). The role of the hippocampus in navigation is memory. Journal of Neurophysiology, 117(4):1785–1796.

Eichenbaum, H., Dudchenko, P., Wood, E., Shapiro, M., and Tanila, H. (1999). The hippocampus, memory, and place cells: Is it spatial memory or a memory space? Neuron, 23(2):209–26.

Fortin, N. J., Agster, K. L., and Eichenbaum, H. B. (2002). Critical role of the hippocampus in memory for sequences of events. Nature Neuroscience, 5(5):458–462.

Franzius, M., Sprekeler, H., and Wiskott, L. (2007). Slowness and Sparseness Lead to Place, Head-Direction, and Spatial-View Cells. PLOS Computational Biology, 3(8):e166.

Hafting, T., Fyhn, M., Molden, S., Moser, M.-B., and Moser, E. I. (2005). Microstructure of a spatial map in the entorhinal cortex. Nature, 436(7052):801–806.

Howard, M. W. and Kahana, M. J. (2002). A Distributed Representation of Temporal Context. Journal of Mathematical Psychology, 46(3):269–299.

Irie, K., Schlag, I., Csordás, R., and Schmidhuber, J. (2021). Going Beyond Linear Transformers with Recurrent Fast Weight Programmers. 2106.06295.

Jacobs, L. F. (2003). The evolution of the cognitive map. Brain, Behavior and Evolution, 62(2):128–139.

Jeffery, K. J. and Burgess, N. (2006). A metric for the cognitive map: Found at last? Trends in Cognitive Sciences, 10(1):1–3.

Jercog, P. E., Ahmadian, Y., Woodruff, C., Deb-Sen, R., Abbott, L. F., and Kandel, E. R. (2019). Heading direction with respect to a reference point modulates place-cell activity. Nature Communications, 10(1):2333.

Leutgeb, J. K., Leutgeb, S., Treves, A., Meyer, R., Barnes, C. A., McNaughton, B. L., Moser, M.-B., and Moser, E. I. (2005). Progressive Transformation of Hippocampal Neuronal Representations in “Morphed” Environments. Neuron, 48(2):345–358.

Lisman, J., Buzsáki, G., Eichenbaum, H., Nadel, L., Rangananth, C., and Redish, A. D. (2017). Viewpoints: How the hippocampus contributes to memory, navigation and cognition. Nature Neuroscience, 20(11):1434–1447.

Markus, E. J., Qin, Y. L., Leonard, B., Skaggs, W. E., McNaughton, B. L., and Barnes, C. A. (1995). Interactions between location and task affect the spatial and directional firing of hippocampal neurons. The Journal of neuroscience : the official journal of the Society for Neuroscience, 15(11):7079–7094.

McInnes, L., Healy, J., and Melville, J. (2020). UMAP: Uniform Manifold Approximation and Projection for Dimension Reduction. 1802.03426.

McNaughton, B. L., Barnes, C. A., and O’Keefe, J. (1983). The contributions of position, direction, and velocity to single unit activity in the hippocampus of freely-moving rats. Experimental Brain Research, 52(1):41–49.

McNaughton, B. L., Battaglia, F. P., Jensen, O., Moser, E. I., and Moser, M.-B. (2006). Path integration and the neural basis of the ‘cognitive map’. Nature Reviews Neuroscience, 7(8):663–678.

Moita, M. A., Rosis, S., Zhou, Y., LeDoux, J. E., and Blair, H. T. (2003). Hippocampal place cells acquire location-specific responses to the conditioned stimulus during auditory fear conditioning. Neuron, 37(3):485–497.

Muller, R. U., Bostock, E., Taube, J. S., and Kubie, J. L. (1994). On the directional firing properties of hippocampal place cells. The Journal of neuroscience : the official journal of the Society for Neuroscience, 14(12):7235–51.

Navawongse, R. and Eichenbaum, H. (2013). Distinct Pathways for Rule-Based Retrieval and Spatial Mapping of Memory Representations in Hippocampal Neurons. Journal of Neuroscience, 33(3):1002– 1013.

O’Keefe, J. (1976). Place units in the hippocampus of the freely moving rat. Experimental Neurology, 51(1):78–109.

O’Keefe, J. and Nadel, L. (1978). The Hippocampus as a Cognitive Map. Clarendon Press, Oxford.

Omer, D. B., Maimon, S. R., Las, L., and Ulanovsky, N. (2018). Social place-cells in the bat hippocampus. Science, 359(6372):218–224.

Parra-Barrero, E., Vijayabaskaran, S., Seabrook, E., Wiskott, L., and Cheng, S. (2023). A map of spatial navigation for neuroscience. Neuroscience & Biobehavioral Reviews, 152:105200.

Quiroga, R. Q., Reddy, L., Kreiman, G., Koch, C., and Fried, I. (2005). Invariant visual representation by single neurons in the human brain. Nature, 435(7045):1102–1107.

Ravassard, P., Kees, A., Willers, B., Ho, D., Aharoni, D. A., Cushman, J., Aghajan, Z. M., and Mehta, M. R. (2013). Multisensory control of hippocampal spatiotemporal selectivity. Science, 340(6138):1342– 1346.

Rolls, E. T. (2010). A computational theory of episodic memory formation in the hippocampus. Behavioural brain research, 215(2):180–196.

Rolls, E. T. (2013). The mechanisms for pattern completion and pattern separation in the hippocampus. Frontiers in Systems Neuroscience, 7:74.

Rubin, A., Yartsev, M. M., and Ulanovsky, N. (2014). Encoding of Head Direction by Hippocampal Place Cells in Bats. Journal of Neuroscience, 34(3):1067–1080.

Samsonovich, A. and McNaughton, B. L. (1997). Path integration and cognitive mapping in a continuous attractor neural network model. The Journal of Neuroscience, 17(15):5900–20.

Sarel, A., Finkelstein, A., Las, L., and Ulanovsky, N. (2017). Vectorial representation of spatial goals in the hippocampus of bats. Science, 355(6321):176–180.

Schlag, I., Munkhdalai, T., and Schmidhuber, J. (2021). Learning Associative Inference Using Fast Weight Memory. 2011.07831.

Schulman, J., Wolski, F., Dhariwal, P., Radford, A., and Klimov, O. (2017). Proximal Policy Optimization Algorithms. 1707.06347.

Scoville, W. B. and Milner, B. (1957). Loss of recent memory after bilateral hippocampal lesions. Journal of Neurology, Neurosurgery & Psychiatry, 20:11–21.

Taube, J. S., Muller, R. U., and Ranck, J. B. (1990). Head-direction cells recorded from the postsubiculum in freely moving rats. I. Description and quantitative analysis. The Journal of Neuroscience: The Official Journal of the Society for Neuroscience, 10(2):420–435.

van Strien, N. M., Cappaert, N. L. M., and Witter, M. P. (2009). The anatomy of memory: An interactive overview of the parahippocampal–hippocampal network. Nature Reviews Neuroscience, 10(4):272–282.

Vijayabaskaran, S. and Cheng, S. (2022). Navigation task and action space drive the emergence of egocentric and allocentric spatial representations. PLOS Computational Biology, 18(10):e1010320.

Vijayabaskaran, S., Zeng, X., Ghazinouri, B., Wiskott, L., and Cheng, S. (2025). A taxonomy of spatial navigation in mammals: Insights from computational modeling. Neuroscience & Biobehavioral Reviews, 176:106282.

Whittington, J. C. R., Muller, T. H., Mark, S., Chen, G., Barry, C., Burgess, N., and Behrens, T. E. J. (2020). The Tolman-Eichenbaum Machine: Unifying Space and Relational Memory through Generalization in the Hippocampal Formation. Cell, 183(5):1249–1263.e23.

Wills, T. J., Lever, C., Cacucci, F., Burgess, N., and O’Keefe, J. (2005). Attractor Dynamics in the Hippocampal Representation of the Local Environment. Science, 308(5723):873–876.

Zeng, X., Diekmann, N., Wiskott, L., and Cheng, S. (2023). Modeling the function of episodic memory in spatial learning. Frontiers in Psychology, 14:1160648.

Zutshi, I., Apostolelli, A., Yang, W., Zheng, Z. S., Dohi, T., Balzani, E., Williams, A. H., Savin, C., and Buzsáki, G. (2025). Hippocampal neuronal activity is aligned with action plans. Nature, 639(8053):153– 161.

